# Widespread male-female expression imbalance of X-linked genes across phrynosomatid lizards

**DOI:** 10.1101/2025.10.06.680020

**Authors:** Matthew D. Hale, Pietro L. H. de Mello, Daniel T. Nondorf, Christopher D. Robinson, Henry B. John-Alder, Christian L. Cox, Robert M. Cox

## Abstract

Classic theory on sex chromosome evolution predicts that selection should restore ancestral diploid expression for hemizygous X-linked genes in males. However, this dosage compensation is often incomplete, leaving X enriched for genes with female-biased expression. In this context, iguanian lizards are noteworthy among vertebrates because several species from separate families appear to exhibit both near-complete dosage compensation and male-female expression balance across their ancient, homologous X chromosomes. We tested for this pattern in a phrynosomatid iguanian, *Sceloporus undulatus* (Eastern Fence Lizard), and instead found that both ancient and more recently sex-linked regions of the X chromosome are enriched for genes with female-biased expression, regardless of age (neonate, maturing, adult) or tissue (brain, liver, muscle). By expanding our analysis across 10 phrynosomatid species spanning 4 genera, we found that male-female expression imbalance on the ancestral region of X is phylogenetically conserved. We also found that an inferred chromosomal rearrangement in the *S. jarrovii* lineage has resulted in the novel acquisition of female-biased expression by a formerly autosomal region. Whereas sex-biased expression of the ancestral region of X is primarily due to females overexpressing X-linked genes relative to autosomal genes, sex-biased expression of these formerly autosomal genes in *S. jarrovii* is primarily due to males underexpressing this putative neo-X region. We conclude that male-female expression imbalance on X is widespread across phrynosomatids, potentially reflecting both overexpression in females for ancestral regions that have evolved dosage compensation and underexpression in males for neo-X regions in which dosage compensation has yet to evolve.

## Introduction

The evolution of sex chromosomes poses a recurring challenge because sex-limited Y or W chromosomes typically degenerate due to the suppression of recombination with formerly homologous X or Z chromosomes, resulting in the loss of functional gene content on Y or W (Charlesworth 2021; Graves 2016; Saunders and Muyle 2024). Consequently, genes on X or Z become hemizygous in the heterogametic sex, potentially disrupting protein stoichiometry with autosomal gene products and thus eliciting negative effects on cell function and organismal fitness (Bachtrog et al. 2011; Birchler et al. 2005; Mank 2009; Shaw and White 2022; Veitia and Potier 2015). Classic theory by Ohno (1967) predicts that this deleterious hemizygous state should favor the evolution of compensatory mechanisms that restore ancestral levels of expression for X- or Z-linked genes in the heterogametic sex (Graves 2016; Gu and Walters 2017). Although such dosage compensation has evolved in many taxa, sex chromosomes are nonetheless often enriched for genes with sex-biased expression (Albritton et al. 2014; Chen et al. 2020a; Furman et al. 2020). While this enrichment could indicate that sex chromosomes are hotspots for genes with sexually antagonistic fitness effects (Allen et al. 2013; Dean and Mank 2014), it is primarily thought to occur because dosage compensation is incomplete and evolves on a gene-by-gene basis (Chen et al. 2020a; Darolti et al. 2019; Furman et al. 2020; Gu and Walters 2017; Mank 2013, 2009; Mullon et al. 2015), or because dosage compensation in the heterogametic sex results in the overexpression of X- or Z-linked genes in the homogametic sex (Allen et al. 2013; Bista et al. 2021; Mank et al. 2011; Prince et al. 2010; Wright and Mank 2012). This underscores a key distinction between “sex-chromosome dosage compensation” (*i.e.,* equal expression of X or Z relative to autosomes in the heterogametic sex) and “male-female expression balance” (*i.e.,* equal expression of X or Z in the heterogametic sex relative to X or Z in the homogametic sex) as two separate phenomena (Bista et al. 2021; Disteche 2012; Gu and Walters 2017; Mank 2009).

Although some species exhibit both dosage compensation and male-female expression balance (Lucchesi and Kuroda 2015; Metzger et al. 2021; Vicoso and Bachtrog 2015), others appear to lack dosage compensation despite male-female balance (Albritton et al. 2014; Gu et al. 2017; Pessia et al. 2014); lack male-female balance despite dosage compensation (Allen et al. 2013; Muyle et al. 2018; Prince et al. 2010; Schultheiß et al. 2015); or lack both dosage compensation and male-female balance (Chen et al. 2014; Itoh et al. 2010; Mank 2013). Therefore, many exceptions to Ohno’s (1967) pioneering theory of dosage compensation have emerged, and complete dosage compensation across the entire X or Z chromosome is now considered the exception rather than the rule (Chandler 2017; Chen et al. 2020a; Furman et al. 2020; Gu and Walters 2017; Mank 2022, 2013, 2009). Overall, dosage compensation appears to be more common in XY than in ZW systems, but the generality of this trend varies across lineages and its potential underlying causes are unclear (Gu and Walters 2017; Mank 2013; Mullon et al. 2015; Vicoso et al. 2013b). One limitation of our current understanding of dosage compensation and male-female expression balance is that it is derived from a relatively limited diversity of taxa (Chen et al. 2020a; Gu and Walters 2017; Kratochvil et al. 2021; Saunders and Muyle 2024). To help address this limitation, we tested for dosage compensation and male-female expression balance across a large reptile lineage with ancestral XY sex-determination.

Squamate reptiles (lizards and snakes) are a promising group for studies of dosage compensation because they have repeatedly evolved both XY and ZW sex determination, vary widely in the extent of sex chromosome degeneration and heteromorphism, and exhibit frequent rearrangements between sex chromosomes and autosomes (Alam et al. 2018; Deakin and Ezaz 2019; Gamble et al. 2015; Giovannotti et al. 2017; Leaché et al. 2016; Lisachov et al. 2021; Pennell et al. 2015). Yet, dosage compensation and male-female expression balance have only been characterized in a few squamate reptiles, primarily due to the historical lack of high-quality genome assemblies (Pinto et al. 2023). In the viperid rattlesnakes *Crotalus viridis* and *Sistrurus miliarius*, dosage compensation is partial and the Z chromosome lacks male-female expression balance (Schield et al. 2019; Vicoso et al. 2013a). Likewise, the varanid lizard *Varanus komodoensis* and the related helodermatid lizard *Heloderma suspectum* share a homologous Z chromosome that lacks both dosage compensation and male-female expression balance (Rovatsos et al. 2019; Webster et al. 2024). Similarly, the corytophanid lizard *Basiliscus vittatus* and the pygopodid lizard *Lialis burtonis* each have different X chromosomes that lack male-female expression balance (Acosta et al. 2019; Nielsen et al. 2019; Rovatsos et al. 2021).

In contrast to the lack of dosage compensation and expression balance that characterizes many squamates, the anolid lizard *Anolis carolinensis* is one of the few vertebrates to exhibit near-complete dosage compensation and male-female expression balance on its X chromosome (Chen et al. 2020a; Marin et al. 2017; Rovatsos and Kratochvíl 2021; Rupp et al. 2017). Dosage compensation and expression balance in *A. carolinensis* involve an epigenetic mechanism for male-specific upregulation of X that is reminiscent of that found in *Drosophila* (Marin et al.2017). Interestingly, recent data from the phrynosomatid lizard *Urosaurus nigricaudus,* which has an X chromosome homologous to that of *Anolis carolinensis*, suggest the presence of both dosage compensation and male-female expression balance (Davalos-Dehullu et al. 2023). This raises the question of whether the dosage compensation and male-female expression balance observed in *Anolis carolinensis* is conserved across phrynosomatids and other iguanian lineages in the large Pleurodonta clade (>1,250 species). With the exception of corytophanids, all families in Pleurodonta (including the anolids and phrynosomatids mentioned above) share an ancestral XY system that arose over 100 Mya and is thus comparable in age to the ancient eutherian and monotreme XY systems and the avian and ophidian ZW systems (Acosta et al. 2019; Altmanova et al. 2018; Davalos-Dehullu et al. 2023; Marin et al. 2017; Nielsen et al. 2019; Rovatsos et al. 2014b). Regions of this ancient iguanian X chromosome exhibit broad synteny across several anolid and phrynosomatid genomes (Davalos-Dehullu et al. 2023; Geneva et al. 2022; Westfall et al. 2021) and hemizygosity in males of several anolid and phrynosomatid species (Geneva et al. 2022; Rovatsos et al. 2014a), as well as both dosage compensation and male-female expression balance in *A. carolinensis* (Chen et al. 2020a; Marin et al. 2017) and potentially also in phrynosomatids such as *Urosaurus nigricaudus* (Davalos-Dehullu et al. 2023).

In this study, we first assessed synteny of the X chromosome between the phrynosomatid lizard *Sceloporus undulatus* and other iguanian genomes, then quantified sex-specific DNA dosage along the *S. undulatus* X chromosome to identify regions of male hemizygosity. Using transcriptomic data from multiple ages and tissues, we identified genes that are consistently sex-biased in their expression in *S. undulatus*, determined their chromosomal locations, and tested for both sex-chromosome dosage compensation and male-female expression balance. We conducted these tests separately for the ancestral region of X that is hemizygous in males and for a small, more recently X-linked region in *S. undulatus* that maps to autosomes in several other phrynosomatids and in *A. carolinensis*. While replicating these tests in a congener, we uncovered the transcriptomic signature of another recently X-linked region in *S. jarrovii*, which we also tested for dosage compensation and male-female expression balance. Using liver transcriptomes from seven *Sceloporus* species and three representatives of other phrynosomatid genera, we characterized broad evolutionary trends in sex-biased gene expression across the putatively homologous region of their ancient iguanian X chromosome, as well as the two more recently X-linked regions identified within *S. undulatus* and *S. jarrovii*. Collectively, these analyses extend our understanding of dosage compensation in squamate reptiles, reveal that male-female expression imbalance is widespread across phrynosomatid lizards, and suggest that male-female expression imbalance can occur in the presence of dosage compensation for ancient X-linked genes, or in the absence of dosage compensation for more recently X-linked genes.

## Results

### The X Chromosome of *S. undulatus* has an Ancient Hemizygous Region and a Neo-X Region

Westfall et al. (2021) identified the tenth largest scaffold in the *S. undulatus* assembly (NC_056531.1, 16.86 Mbp, hereafter chromosome 10) as the predicted X chromosome. Our analyses confirm that the majority (15.10 Mbp) of *S. undulatus* chromosome 10 is syntenic with the ancient X chromosome in other iguanian genomes, including the more distantly related anolids *Anolis carolinensis* and *A. sagrei* and the more closely related phrynosomatids *Phrynosoma platyrhinos* and *Urosaurus nigricaudus* (Fig. 1; also see Davalos-Dehullu et al. (2023)). In addition, we found that a small (1.76 Mbp), distal region of *S. undulatus* chromosome 10 is syntenic with a portion of a putatively autosomal microchromosome in *Phrynosoma platyrhinos* (Fig. 1A), suggesting that this is a more recently X-linked region in *S. undulatus* (*i.e*., a neo-X region). Targeted BLAST searches of predicted mRNA sequences from this distal region confirmed that individual genes located before 15.10 Mbp on chromosome 10 in *S. undulatus* typically map to X in all other iguanians (Fig. 1B), whereas genes located after 15.10 Mbp only map to X in closely related *S. tristichus* (formerly considered a subspecies of *S. undulatus*) and distantly related *A. sagrei* (due to an independent X-autosome fusion within *Anolis*, see Geneva et al. (2022)). In other phrynosomatids (*U. nigricaudus, P. platyrhinos*) and other distantly related anolids (*A. carolinensis*), genes located after 15.10 Mbp on the *S. undulatus* X chromosome map to putative autosomes or unplaced scaffolds (Fig. 1B).

**Figure 1.**
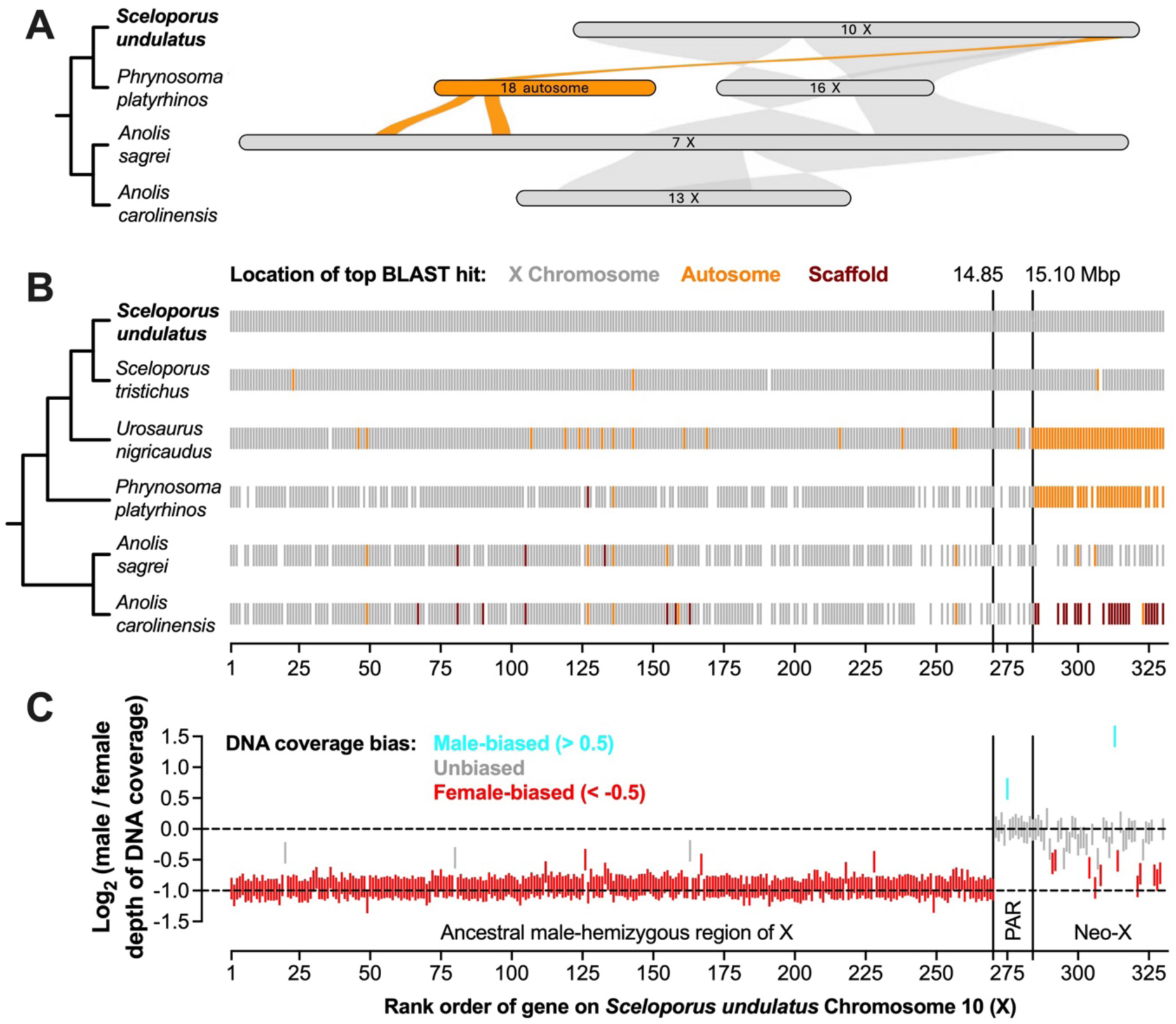
(A) Riparian plot illustrating synteny of *S. undulatus* chromosome 10 (X) with three other iguanian genomes. Only chromosome-level scaffolds with syntenic blocks that map to *S. undulatus* chromosome 10 are shown. **(B)** Chromosomal locations of genes on *S. undulatus* chromosome 10 in five other iguanian genomes, based on top BLAST hits using the longest mRNA isoform for each gene as a query. **(C)** Log_2_ (male/female) depth of DNA sequencing coverage in *S. undulatus* for each gene on chromosome 10, with the dashed line at -1 signifying expected dosage imbalance due to hemizygosity in males. Vertical lines in **B** and **C** indicate inferred boundaries between the ancestral male-hemizygous region, ancestral psuedoautosomal region (PAR), and neo-X region in *S. undulatus* and *S. tristichus*.

Analysis of published whole-genome sequences from 18 female and 11 male *S. undulatus* specimens (Assis et al. 2025) revealed that, as expected, log_2_ (male/female) DNA coverage is centered at 0 (equal dosage) across all chromosomes except chromosome 10 (Table S2; Fig. S1), which exhibits values near -1 along most of its length (Fig. 1C). This two-fold higher DNA coverage in females indicates that males are hemizygous for most of chromosome 10, confirming its identify as the X chromosome. However, chromosome 10 exhibits a distinct shift toward values centered on 0 beginning at approximately 14.85 Mbp, slightly before the start of the putative neo-X region at 15.10 Mbp (Fig. 1C). Whereas most genes between 14.85 and 15.10 Mbp have DNA coverage ratios near 0, genes located after 15.10 Mbp exhibit a range of values between 0 and -1, suggesting that the homologous neo-Y region may be in the process of degeneration (Fig. 1C). In our subsequent analyses of dosage compensation (below), we excluded the 15 genes located between 14.85 and 15.10 Mbp of chromosome 10 because these genes are ancestrally X-linked (Fig. 1B) but exhibit no evidence of a sex difference in dosage (Fig. 1C), suggesting that they may reside on a pseudoautosomal region (PAR) that is still recombining with its Y homolog.

### The *S. undulatus* Genome Contains Many Putatively X- and Y-Linked Scaffolds

Although the *S. undulatus* genome was sequenced from male tissue, the current assembly does not include a formally characterized Y chromosome (Westfall et al. 2021). However, our analyses of unplaced scaffolds revealed 45 annotated genes with mean log_2_ (male/female) DNA coverage values greater than 0.5 (i.e., a male bias in depth of coverage; Table S2). We hypothesize that many of these scaffolds, which each contain a single gene and collectively comprise only 0.2 Mbp, may correspond to Y-linked regions of the S*. undulatus* genome. We also found 116 annotated genes on unplaced scaffolds with mean log_2_ (male/female) DNA coverage values less than -0.5, potentially corresponding to unassembled regions of the X chromosome (Table S2). Filtering to exclude genes with low coverage (mean < 10 in both sexes) left 44 of these potentially Y-linked genes and 65 of these potentially X-linked genes on unassembled scaffolds (Fig. S1).

### The *S. undulatus* Transcriptome Contains Many Consistently Sex-Biased Genes

To characterize male-female expression imbalance in *S. undulatus* without any *a priori* assumptions about genomic location, we conducted genome-wide differential gene expression analyses to identify genes that are consistently sex-biased in their expression across three ages (neonate, maturing, adult) in three distinct tissues (liver, brain, muscle). We reasoned that such consistency is more likely to reflect sex differences in gene dosage than in gene regulation, which is likely to be both age- and tissue-specific. We detected cohorts of significantly differentially expressed genes (DEGs; *P* < 0.05 after correction for false discovery rate) that are consistently female-biased across all ages in the liver (*n* = 32 genes; Fig. 2A-C), brain (*n* = 48; Fig. S2A-C), and muscle (*n* = 11; Fig. S3A-C). We also detected cohorts of DEGs that are consistently male-biased across all ages in the liver (*n* = 27; Fig. 2A-C), brain (*n* = 34; Fig. S2A-C), and muscle (*n* = 19; Fig. S3A-C). These consistently sex-biased DEGs typically exhibit the same patterns of sex bias across tissues (Fig. S4A-I), and many are identified as consistently sex-biased DEGs in more than one tissue (Fig. S4J-K). Consistently female-biased DEGs fall within a relatively narrow range of log_2_ FC values corresponding to approximately 1.5 to 2-fold higher expression in females, while consistently male-biased DEGs exhibit a wider range of more extreme log_2_ FC values (Figs. S2-S5). Consistently female-biased DEGs are expressed in both sexes, as expected if they are located on the X chromosome, whereas many consistently male-biased DEGs are expressed at low or undetectable levels in females (Fig. S5), as expected if they are located on the Y chromosome.

**Figure 2.**
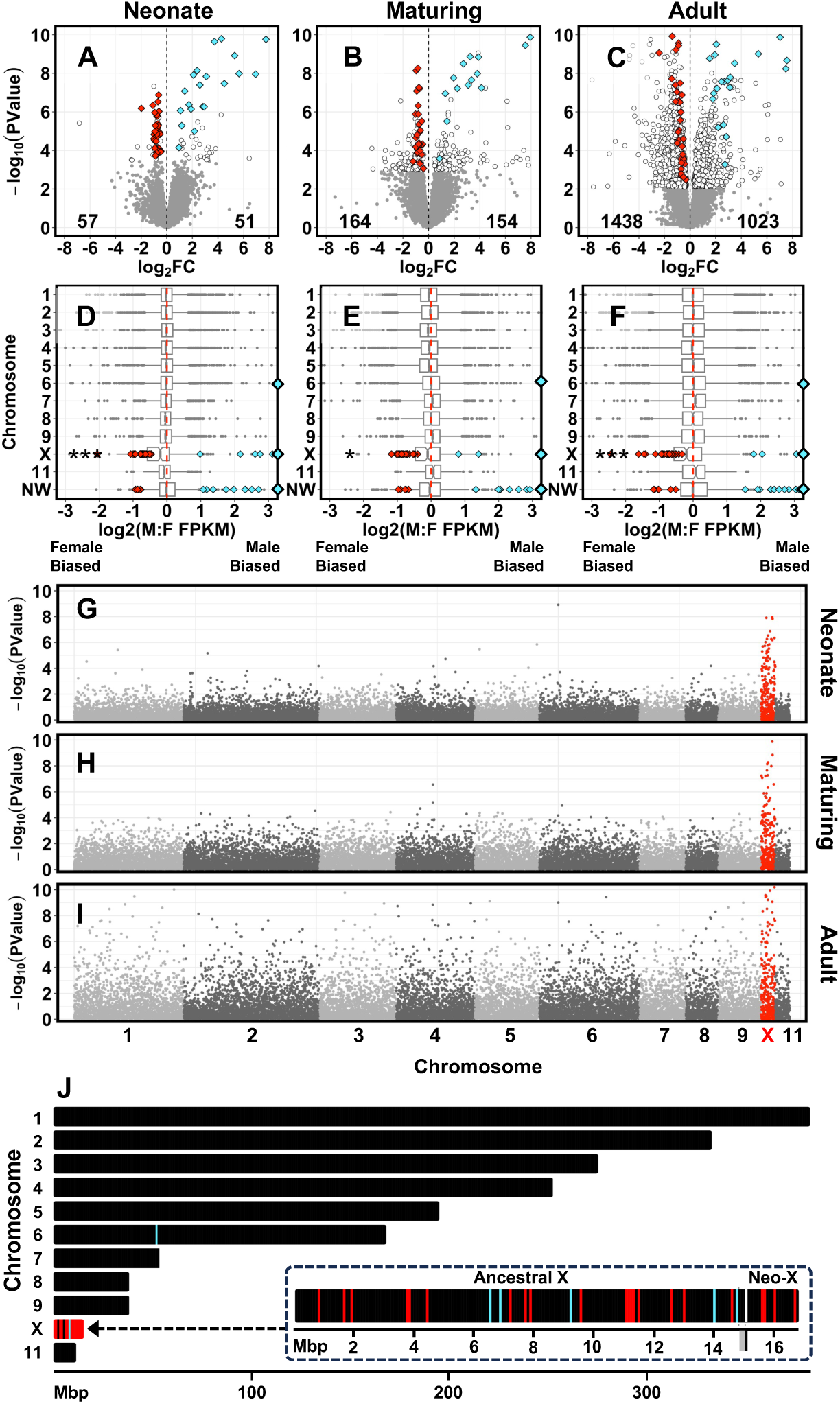
(A-C) Volcano plots depicting the magnitude (log_2_ fold change) and significance (-log_10_ *P* value) of sex-biased expression across 15,587 genes in liver of *S. undulatus* across three ages. Genes with significantly (FDR < 0.05) sex-biased expression (DEGs) are indicated in white, with the number of female-biased and male-biased DEGs indicated in the bottom corners of each panel. DEGs with consistently sex-biased expression across all ages are shown as red (female-biased) or blue (male-biased) diamonds. Grey points are genes that are not significantly sex-biased. **(D-F)** Boxplots depicting the median (line), interquartile range (box) and the smallest or largest values within 1.5 x IQR (whiskers) for sex-biased expression (log_2_ male:female ratio of fragments per kilobase per million mapped reads, FPKM) from each of the 11 largest scaffolds and pooled genes from unplaced scaffolds (NW) in the *S. undulatus* assembly. The 10^th^ largest scaffold is the X chromosome. Notches in the center of each boxplot indicate the 95% confidence interval of the median. Diamonds indicate consistently female-biased (red) and male-biased (blue) DEGs in each tissue, which map predominantly to chromosome 10 (X) and unplaced scaffolds (NW). Diamonds on plot borders indicate DEGs beyond x-axis limits. Asterisks indicate when median chromosome expression deviates significantly from balanced male:female expression using Wilcoxon rank sum testing (mu = 0) with correction for multiple tests (* *P* < 0.05 without correction for multiple tests; *\*\* P* < 0.017, the critical alpha after Bonferroni correction; *** *P* < 0.001). **(G-I)** Manhattan plots of -log_10_ *P* values for sex-biased expression after ordering genes by chromosome and start position. All genes on X are shown in red. **(J)** Genomic locations of consistently sex-biased DEGs (unplaced scaffolds not shown). Enlarged inset shows that these genes occur on both the ancestral region of X that is hemizygous in males and on the putative neo-X region beyond the large tick mark at 15.1 Mbp. Gray shading immediately before this large tick indicates the putative psuedoautosomal region.

### The X Chromosome of *S. undulatus* is a Hotspot for Female-Biased Gene Expression

The majority (45 of 60) of the consistently female-biased DEGs we identified are located on the X chromosome (Fig. 2D-F; Figs. S2-S3). The remainder (15 of 60) are located on small, unplaced scaffolds, many (9 of 15) of which have log_2_ (male/female) DNA coverage values below -0.5, suggesting that they are regions of X that were orphaned during assembly (Table S2). Genes across the X chromosome collectively exhibit a significant female bias in their median expression at 2 of 3 ages in liver (Fig. 2D-F), 3 of 3 ages in brain (Fig. S2D-F) and 1 of 3 ages in muscle (Fig. S3D-F). We found identical results when restricting this comparison to genes in the proximal 14.85 Mbp, ancestrally X-linked region. Plotting sex-biased expression as a function of chromosomal location across the 11 largest scaffolds in the *S. undulatus* genome reveals a single peak of elevated sex bias on the X chromosome that is evident across all ages and tissues (Fig. 2G-I; Figs. S2-S3). Consistently female-biased genes are widely distributed across the entire length of the X chromosome and are interspersed with many unbiased genes and several consistently male-biased genes (Fig. 2J; Fig. S6). Of the 6 consistently female-biased DEGs located in the hypothesized neo-X region, 5 exhibit a negative log_2_ (male/female) DNA coverage value below -0.25, suggesting that their gametologs on the neo-Y chromosome have degenerated or diverged. Whereas both the proximal 14.85 Mbp (ancestrally X-linked region) and the distal 1.76 Mbp (hypothesized neo-X region) of the X chromosome contain consistently female-biased DEGs and exhibit an overall tendency toward female-biased expression (Fig. S6), the intermediate 0.25 Mbp (hypothesized PAR) exhibits virtually no sex-biased expression at any age or in any tissue (Fig. S6). Enrichment of the X chromosome with female-biased genes was recapitulated when we mapped *S. undulatus* reads to the genome of closely related *S. tristichus* (Bedoya and Leaché 2021), which was sequenced from female tissue and therefore lacks any Y-linked sequence in its assembly (Figs. S7-S8, see Supplementary Results).

### Consistently Male-Biased Genes Map to X and to Putative Y-linked Scaffolds in *S. undulatus*

The majority (31 of 45) of the consistently male-biased DEGs that we identified are located on small, unplaced scaffolds in the *S. undulatus* assembly (Fig. 2D-F; Figs. S2-S3), most of which (22 of 31) we hypothesized to be Y-linked based on log_2_ (male/female) DNA coverage values that are above 0.5 (Table S2). One of these male-biased DEGs (*LOC121918292*, an *RPL6*-like gene) had no detectable DNA reads from females. *RPL6* is Y-linked in *Anolis carolinensis* (Marin et al. 2017) and is the putative sex-determining locus in *A. sagrei* (Motley 2025). A subset (10 of 45) of consistently male-biased DEGs are located on the X chromosome.

Although this could potentially occur if some Y-linked regions co-assembled with X (Westfall *et al*., 2021), all of these male-biased genes on the X chromosome exhibit log_2_ (male/female) DNA coverage values near -1 (Fig. 1C), indicating that they are hemizygous in males. A few (4 of 45) consistently male-biased DEGs are located on putative autosomes (Figs. 2D-F; S3D-F).

### The X Chromosome of *S. undulatus* is Overexpressed Relative to Autosomes in Females

To determine whether male-female expression imbalance on the *S. undulatus* X chromosome is driven by the lack of dosage compensation in males versus overcompensation in females, we compared the expression of X-linked genes to the expression of autosomal genes separately within each sex. We did this by randomly sampling 1000 contiguous regions of the autosomal genome with the same number of expressed genes as the corresponding regions of X to generate null distributions of median autosomal expression values for each sex, age, and tissue (Fig. 3). In *S. undulatus* males, the median expression of ancestrally X-linked genes on the proximal 14.85 Mbp region of the X chromosome is statistically indistinguishable from the sampled autosomal distributions for liver and trends toward the high ends of the sampled autosomal distributions for brain and muscle (Fig. 3A). In females, the median expression of these same ancestrally X-linked genes trends toward the high ends of the simulated autosomal distributions for liver and muscle and significantly exceeds the simulated autosomal distributions for brain (Fig. 3A). We observed similar patterns for more recently X-linked genes on the distal, 1.76-Mbp region of the X chromosome (Fig. 3B). Collectively, these results indicate that female-biased expression across the X chromosome is associated with elevated expression of X-linked genes in females, rather than reduced expression of X-linked genes in males.

**Figure 3.**
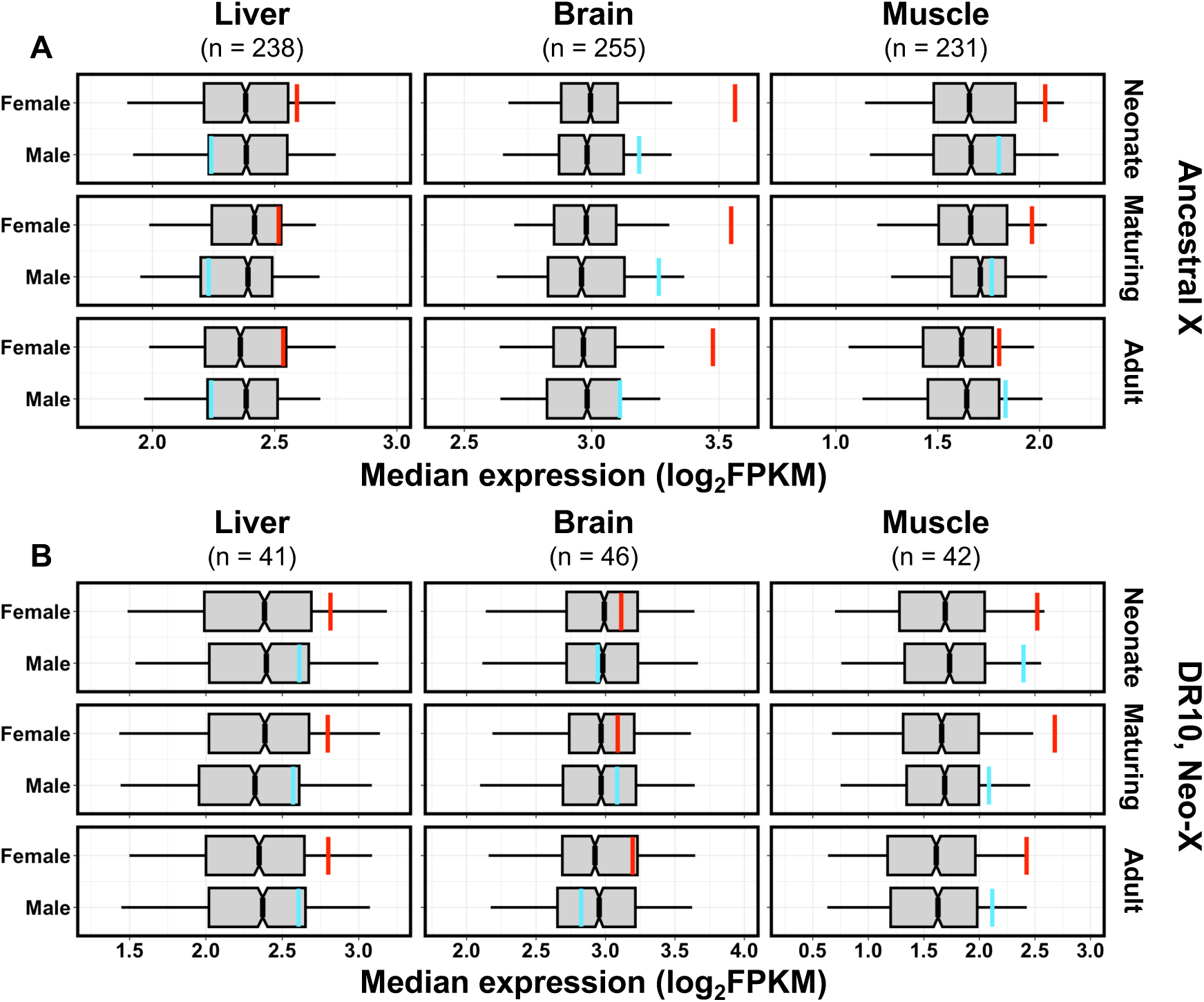
Comparisons of median expression on the X chromosome versus that of autosomes for each sex across three tissues and three ages in *S. undulatus*, shown separately for (**A**) the ancestrally X-linked 14.85-Mbp region of chromosome 10, and (**B**) the distal 1.76 Mbp region (DR10) that is more recently X-linked. Boxplots depict the median (line), IQR (box), and 5^th^ and 95^th^ quantiles (whiskers) for median expression estimates from 1000 randomly-sampled blocks of *n* contiguous autosomal genes (*n* = number of expressed genes in the corresponding region of the X chromosome in that tissue). Colored vertical lines (red = female; blue = male) indicate the median expression of all genes on X for each combination of sex, age, and tissue. Colored lines which fall beyond whiskers indicate that median expression of genes on the X chromosome differs significantly from the sampled distribution for autosomal genes.

### Consistently Sex-Biased Genes Identify a Putative Neo-X Region in *S. jarrovii*

To explore the phylogenetic conservation of patterns identified in *S. undulatus,* we also tested for consistently sex-biased genes using liver transcriptomes from *S. jarrovii* that were sampled at three ages (neonate, maturing, adult). We detected 63 DEGs in *S. jarrovii* liver that are consistently female-biased across all ages (Fig. 4A-C), as well as 12 DEGs that are consistently male-biased across all ages (Fig. 4A-C). These consistently female-biased DEGs fall within a relatively narrow range of log_2_ FC values that corresponds to approximately 2-fold higher expression in females than in males, consistent with X-linkage, whereas most of the consistently male-biased DEGs are expressed at low or undetectable levels in females, consistent with Y-linkage (Fig. S10). Many (20 of 63) of the consistently female-biased DEGs in *S. jarrovii* map to the X chromosome in *S. undulatus* (Fig. 4D-F), and the median expression of all genes that map to the *S. undulatus* X chromosome is significantly female-biased at all ages in *S. jarrovii* (Fig. 4D-F). None of the consistently sex-biased genes in *S. jarrovii* map to the distal 1.76-Mbp region of the *S. undulatus* X chromosome (Fig. 4J), and this region exhibits no overall female bias in its expression in *S. jarrovii* (Fig. S11), as expected given that genes in this region map to autosomes in all phrynosomatids other than *S. undulatus* and *S. tristichus* (Fig. 1).

**Figure 4.**
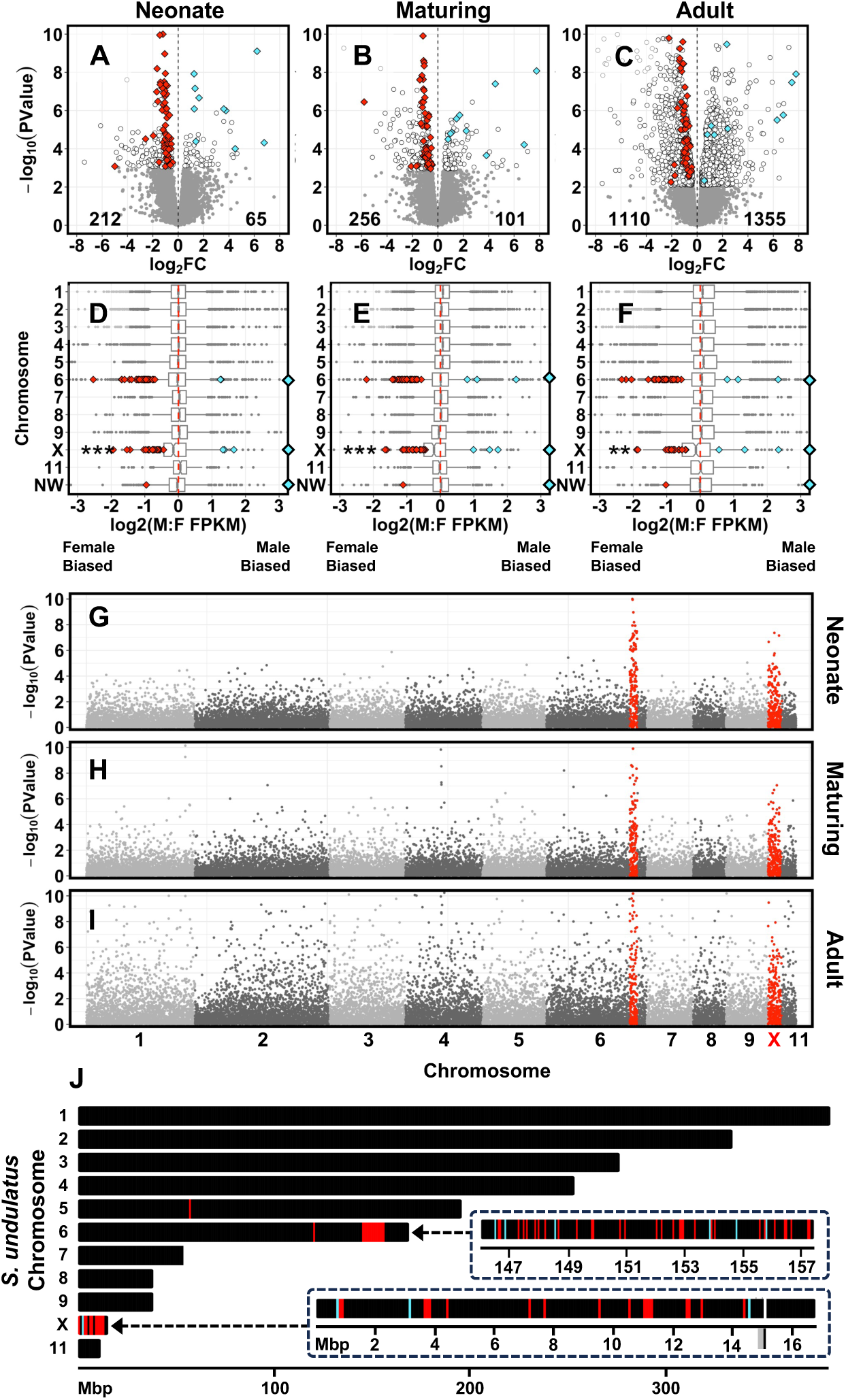
(A-C) Volcano plots depicting the magnitude (log_2_ fold change) and significance (-log_10_ *P* value) of sex-biased expression across 15,493 genes in liver of *S. jarrovii.* Genes with significantly (FDR < 0.05) sex-biased expression (DEGs) are indicated in white, with the number of female-biased and male-biased DEGs indicated in the bottom corners of each panel. DEGs with consistently sex-biased expression across all ages are shown as red (female-biased) or blue (male-biased) diamonds. Grey points depict genes that are not significantly sex-biased. (**D-F**) Boxplots depicting the median (line), interquartile range (box) and the smallest or largest values within 1.5 x IQR (whiskers) for sex-biased expression (log_2_ male:female ratio of fragments per kilobase per million mapped reads, FPKM) from each of the 11 largest scaffolds and pooled genes from unplaced scaffolds (NW) in the *S. undulatus* genome. Diamonds indicate consistently female-biased (red) and male-biased (blue) DEGs, which map predominantly to chromosome 6, chromosome 10 (X), and unplaced scaffolds (NW). Diamonds on plot borders indicate genes beyond x-axis limits. Asterisks indicate that median chromosome expression deviates significantly from balanced male:female expression using Wilcoxon rank sum testing (mu = 0) with correction for multiple tests (* *P* < 0.05 without correction for multiple tests; *\*\* P* < 0.017, the critical alpha after Bonferroni correction; *** *P* < 0.001). (**G-I**) Manhattan plots depicting - log_10_ *P* values for sex-biased expression after ordering genes by *S. undulatus* chromosome and start position. All genes on X and within an ∼11 Mpb distal region of chromosome 6 (DR6) are shown in red. (**J**) Locations of consistently sex-biased DEGs in *S. jarrovii* when mapped to the *S. undulatus* genome. All consistently sex-biased DEGs map to the ancestral X chromosome (the region of *S. undulatus* chromosome 10 prior to the large tick at 15.1 Mbp in the lower inset) or the distal ∼11 Mbp region of chromosome 6 (DR6, upper inset), suggesting that this region of *S. undulatus* chromosome 6 is X-linked in *S. jarrovii*.

Unlike in *S. undulatus*, most of the consistently female-biased DEGs (41 of 63) and half of the consistently male-biased DEGs (6 of 12) that we detected in *S. jarrovii* transcriptomes map to the sixth largest scaffold in the *S. undulatus* assembly (NC_056527.1, hereafter chromosome 6), which is an autosome in *S. undulatus* (Fig. 4D-F). Moreover, nearly all (46 of 47) of these consistently sex-biased DEGs fall within a relatively small, 11-Mbp distal region of chromosome 6 (hereafter DR6; Fig. 4J) that exhibits a sharp peak of sex-biased expression similar to that observed on the X chromosome (Fig. 4G-I). Genes on chromosome 6 that fall outside of DR6 exhibit no overall tendency toward sex-biased expression in either *S. jarrovii* or *S. undulatus* (Fig. 5). However, the 156 genes located within DR6 are significantly female-biased in *S. jarrovii*, even when excluding the subset of 46 consistently sex-biased DEGs that we first used to identify this region (Fig. 5B). These 156 genes within DR6, including the 46 consistently sex-biased DEGs from *S. jarrovii*, do not exhibit any overall sex bias in DNA coverage (Fig. S1) or expression (Fig. 5B) in *S. undulatus*. Collectively, genes within DR6 exhibit expression profiles similar to the X chromosome in *S. jarrovii,* but similar to autosomes in *S. undulatus* (Fig. 5). Our expression data therefore suggest that the *S. jarrovii* X chromosome shares an ancestral 14.85-Mbp region with *S. undulatus,* lacks the 1.76-Mbp region that is more recently X-linked in *S. undulatus* (DR10), but includes an 11-Mbp region that is autosomal in *S. undulatus* and more recently X-linked in *S. jarrovii* (DR6).

**Figure 5.**
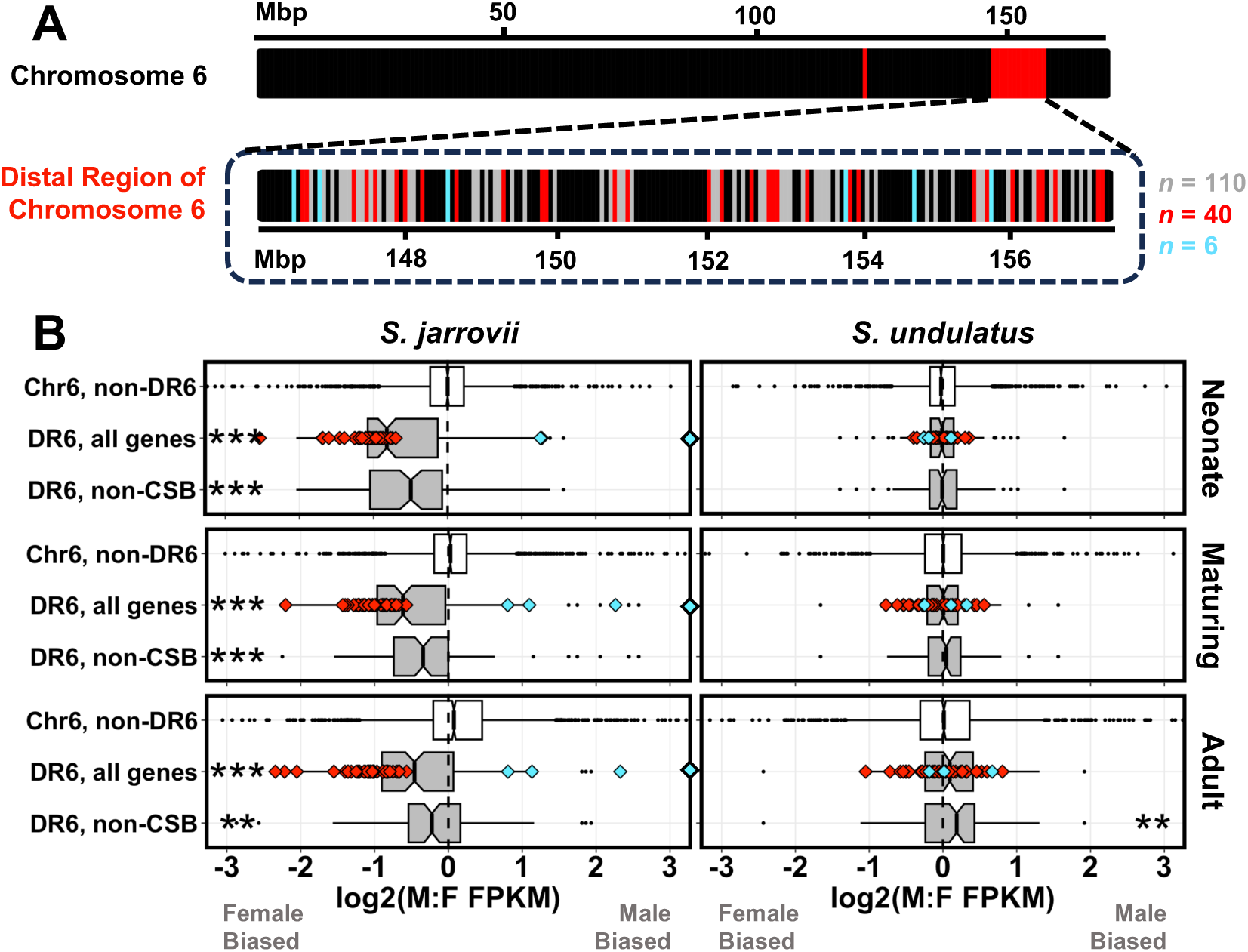
(**A**) Location of consistently female-biased (red), male-biased (blue), and unbiased genes (gray) in *S. jarrovii* that map to the distal region of chromosome 6 (DR6) in *S. undulatus.* Bars are not drawn to scale for gene length. (**B**) Boxplots depicting median (line), interquartile range (box), and 1.5 x IQR (whiskers) for sex-biased expression (log_2_ male:female ratio of fragments per kilobase per million mapped reads, FPKM) in liver of *S. jarrovii* and *S. undulatus* at three ages. Boxplots are show for all genes on chromosome 6 except those within DR6 (“Chr 6, non-DR6”), for all genes within DR6 (“DR6, all genes”), and for all genes within DR6 excluding consistently sex-biased DEGs (“DR6, non-CSB”). Diamonds depict consistently sex-biased DEGs with those on plot borders indicating sex bias beyond axis limits. Asterisks indicate when median expression differs from zero using Wilcoxon signed-rank tests with Bonferroni correction for 2 tests per age (* 0.025 > *P* > 0.001; ** 0.001 > *P* > 0.0001; *** *P* < 0.0001).

### The Putative Neo-X Region in *S. jarrovii* Lacks Dosage Compensation in Males

Similar to our analyses for *S. undulatus*, we compared the median expression of putatively X-linked genes in *S. jarrovii* (*i.e.,* genes on the ancestrally X-linked 14.85-Mbp region of *S. undulatus* chromosome 10 and the recently X-linked 11-Mbp DR6 region of *S. undulatus* chromosome 6) to simulated null distributions of median autosomal expression in males and females by randomly sampling 1000 contiguous regions of the autosomal genome with the same number of expressed genes as the corresponding X-linked regions. In *S. jarrovii* males, median levels of expression for genes mapping to the ancestrally X-linked region of *S. undulatus* chromosome 10 fall within the simulated distributions for autosomal genes, but lower than the median (Fig. S12A). In *S. jarrovii* females, median levels of expression for these same genes are toward the high end of sampled distributions for autosomal genes (Fig. S12A). Therefore, the overall pattern of female-biased expression for ancestrally X-linked genes in *S. jarrovii* reflects a combination moderate underexpression in males and moderate overexpression in females (Fig. S12A). In both sexes of *S. jarrovii*, the median expression of genes that map to DR10 in *S. undulatus* is near the center of the simulated autosomal distribution (Fig. S12B), consistent with the interpretation that this neo-X region in *S. undulatus* is not X-linked in *S. jarrovii*.

By contrast, the median expression of genes within DR6 is significantly reduced in *S. jarrovii* males when compared to sampled null distributions for autosomal genes for maturing and adult ages and nearly so for neonates (Fig. 6A). In *S. jarrovii* females, the median expression of genes within DR6 is similar to the simulated distributions for autosomal genes at all ages (Fig. 6A). In *S. undulatus*, the median expression of genes within DR6 is similar to the simulated distribtions for autosomal genes in both sexes (Fig. 6A). Therefore, DR6 in *S. jarrovii* exhibits the predicted transcriptomic profile for X-linked genes that lack dosage compensation in males, whereas DR6 in *S. undulatus* is expressed at levels comparable to other autosomal genes in both sexes. Paired, inter-species comparisons of *S. jarrovii* and *S. undulatus* homologs on DR6 corroborate this interpretation. Homologous genes on DR6 are expressed at comparable levels in females of each species (paired *t*-tests within each age, all *P* > 0.16; Fig. 6B), but at significantly lower levels in *S. jarrovii* males than in *S. undulatus* males (all *P* < 0.001; Fig. 6B).

**Figure 6.**
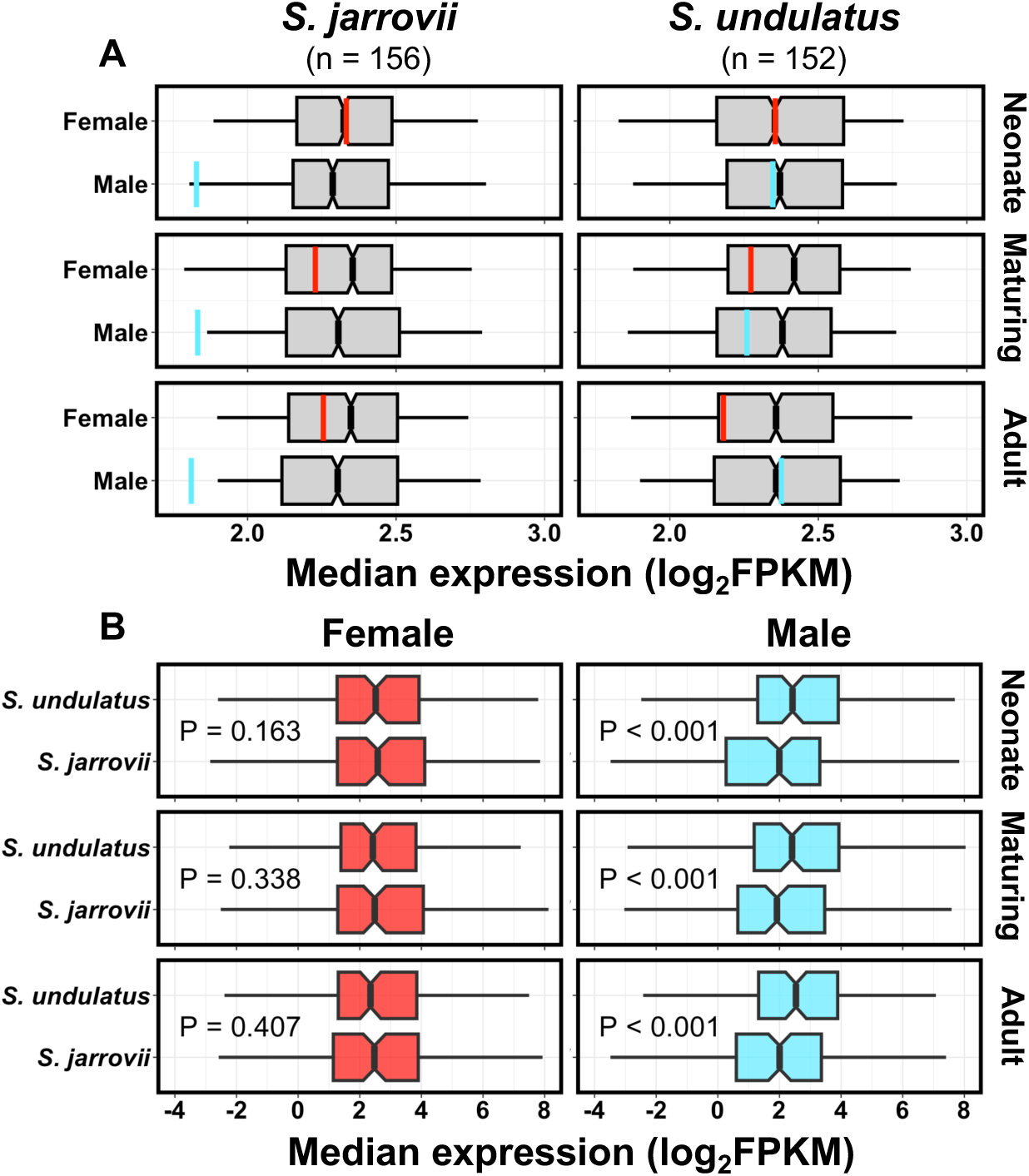
(**A**) Median expression of genes in the distal region of *S. undulatus* chromosome 6 (DR6) versus a sampled distribution of median autosomal expression for each sex across three ages in *S. jarrovii* and *S. undulatus*. Boxplots depict the median (line), intraquartile range (box), and 5^th^ and 95^th^ quantiles for median expression values from 1000 randomly sampled blocks of *n* contiguous autosomal genes (*n* = number of expressed genes within DR6 in each species). Vertical red (female) and blue (male) lines indicate median expression of all genes within DR6. (**B**) Comparisons (paired *t-*test by gene) of expression for all genes on DR6 between *S. jarrovii* and *S. undulatus,* conducted separately by sex. Boxplots depict median (line), intraquartile range (box), and 2.5^th^ and 97.5^th^ quantiles for expression of 152 genes present on DR6 in both species.

### Male-Female Expression Imbalance of the Ancestral X is Conserved Across Phrynosomatids

To characterize broad evolutionary trends in the expression of sex-linked genes, we combined our expression data from adult livers in *S. undulatus* and *S. jarrovii* with similar RNA-seq data (mapped to the *S. undulatus* genome) from adult livers in 5 other *Sceloporus* species, plus representatives of 3 other phrynosomatid genera (Table S1). Across Phrynosomatidae, we observed female-biased expression of genes mapping to the ancestrally X-linked 14.85-Mbp region of *S. undulatus* chromosome 10, suggesting that this signature of male-female expression imbalance is the ancestral state for this family (Fig. 7A). Median log_2_ (male/female) expression ratios across this putatively homologous region of the X chromosome are significantly female-biased in adult liver from 6 of 7 *Sceloporus* species, as well as from all 3 phrynosomatid species representing other genera (Fig. 7A). Moreover, each of the DEGs from this ancestral region of chromosome 10 that are consistently female-biased across ages in *S. undulatus* are nearly always also female-biased in other phrynosomatid species (Fig. 7A). Likewise, the few DEGs from this ancestral region of chromosome 10 that are consistently male-biased across ages in *S. undulatus* also tend to exhibit male-biased expression in other species, although this conservation is somewhat weaker (Fig. 7A).

**Figure 7.**
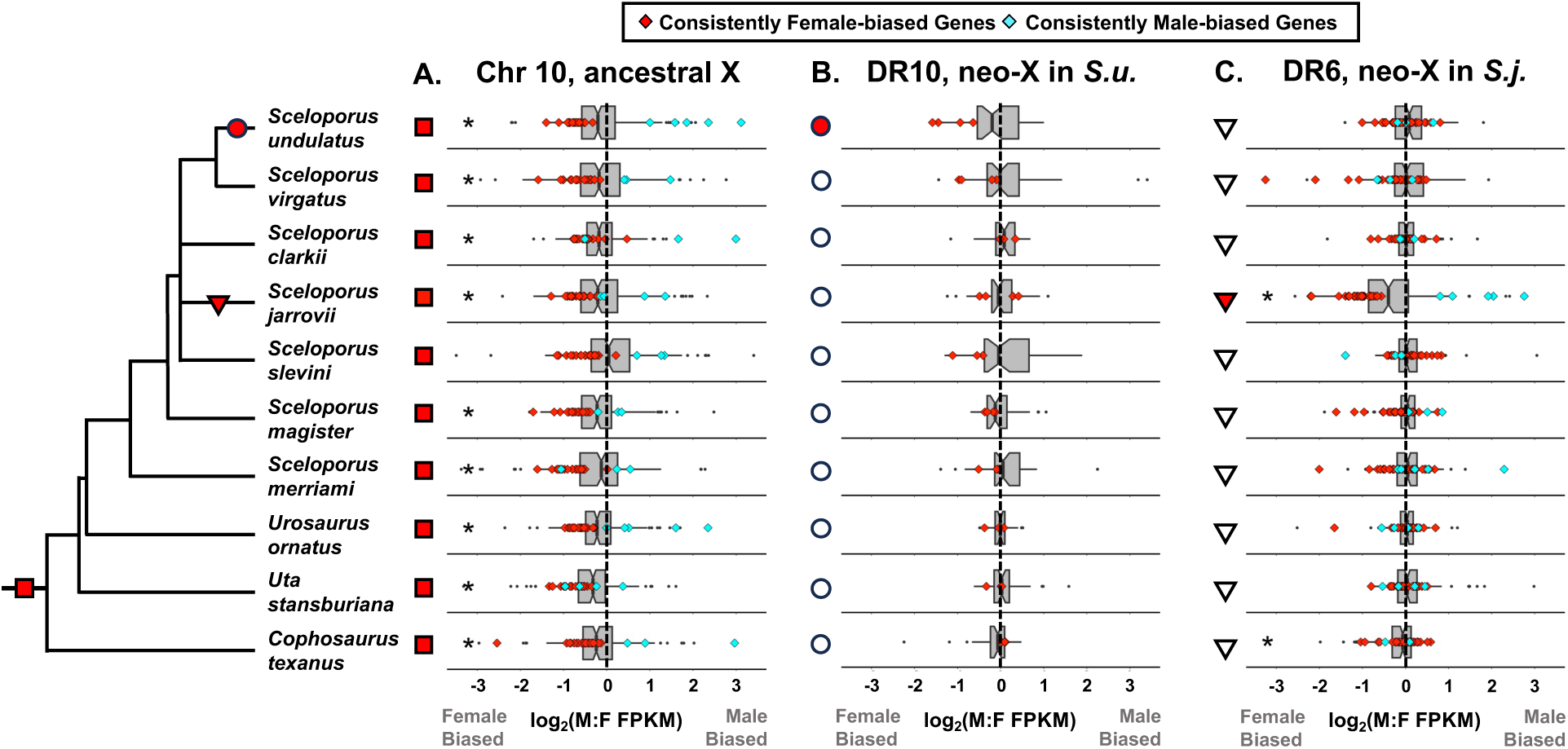
Evolutionary patterns in sex-biased expression of (**A**) the 14.85-Mbp region of *S. undulatus* chromosome 10 that is hemizygous in males (putative ancestral X), (**B**) the distal 1.76-Mbp region of chromosome 10 (DR10, putative neo-X region in *S. undulatus*), and (**C**) the distal 11-Mbp region of chromosome 6 (DR6, putative neo-X region in *S. jarrovii*) based on adult liver transcriptomes from 10 phrynosomatid species mapped to the *S. undulatus* assembly. Boxplots depict the median (line), intraquartile range (box), and 1.5 x IQR (whiskers) of log_2_ male:female expression (FPKM) for all expressed genes within each region. Diamonds indicate consistently female-biased (red) and male-biased (blue) genes in transcriptomes from *S. undulatus* (**A-B**) or *S. jarrovii* (**C**). Notches in boxplots indicate 95% confidence intervals of the median. Asterisks denote whether median expression ratios differ significantly from zero (Wilcoxon signed-rank test, Bonferroni-adjusted alpha = 0.005). Red symbols to the left of each boxplot indicate when consistently female-biased DEGs from *S. undulatus* (**A-B**) or *S. jarrovii* (**C**) differ from zero in each species (Wilcoxon signed-rank test, Bonferroni-adjusted alpha = 0.005) and are used to map ancestral (X) and derived (neo-X) male-female expression imbalances onto the phylogeny.

### Male-Female Expression Imbalance of Neo-X Regions is Restricted to Individual Lineages

In contrast to the phylogenetically conserved patterns of sex-biased expression for genes on the ancestrally X-linked region of *S. undulatus* chromosome 10 (Fig. 7A), only *S. undulatus* exhibits female-biased expression of genes on its putative neo-X region comprising the distal 1.76 Mbp of chromosome 10 (DR10, Fig. 7B). Moreover, the few genes from DR10 that are consistently female-biased DEGs across ages in *S. undulatus* exhibit no overall tendency for female-biased expression in other species (Fig. 7B). Similarly, only *S. jarrovii* exhibits female-biased expression for genes on its putative neo-X region comprising the 11-Mbp distal portion of chromosome 6 in *S. undulatus* (DR6, Fig. 7C). Likewise, genes from DR6 that are consistently either female- or male-biased DEGs across ages in *S. jarrovii* exhibit no clear conservation of these sex-biased expression patterns across other phrynosomatid species (Fig. 7C). Overall, these evolutionary patterns of sex-biased gene expression indicate that the 1.76-Mbp distal region of chromosome 10 is a recently X-linked region in *S. undulatus* (and closely related species such as *S. tristichus*, Fig. 1B), and that the 11-Mbp distal region of chromosome 6 in *S. undulatus* has also recently become sex-linked in the lineage represented here by *S. jarrovii* (Fig. 7C).

## Discussion

Recent research on sex chromosomes has emphasized the distinction between dosage compensation (of X or Z with autosomes in the heterogametic sex) and expression balance (of X or Z between the sexes) while noting that associations between the two can differ across lineages (Bista et al. 2021; Gu and Walters 2017; Saunders and Muyle 2024). In *Sceloporus undulatus,* male-female expression imbalance across the ancestral, male-hemizygous region of X appears to be associated with the overexpression of this region (relative to autosomes) in females, rather than its underexpression in males. Although this “Type IV” profile (*i.e.,* dosage compensation without male-female expression balance, following Gu & Walters 2017) has been observed in a variety of taxa (Allen et al. 2013; Muyle et al. 2018; Prince et al. 2010; Schultheiß et al. 2015), several subsequent studies have attributed it to artifacts from microarray data, the inclusion of gonadal tissue, or the lack of appropriate tests for dosage compensation (Gu and Walters 2017; Saunders and Muyle 2024). These first two concerns are not relevant to our study, but it has been proposed that tests for dosage compensation should ideally compare expression between sex-linked genes and their orthologs still in the ancestral autosomal state (Bista et al. 2021; Gu and Walters 2017). Due to the considerable age (>100 Mya) of the iguanian X chromosome that is ancestral to Pleurodonta (Acosta et al. 2019; Davalos-Dehullu et al. 2023; Marin et al. 2017), this would require comparisons across distantly related lineages in which transcriptomic profiles may have diverged considerably over millenia. Instead, by comparing X to null distributions sampled from contiguous autosomal blocks within the same genome (and the same biological replicates), we provide an alternative approach that suggests an overall Type IV expression profile for the ancestral region of the X chromosome in *S. undulatus*. We emphasize that this is a summary categorization across hundreds of X-linked genes that vary considerably in their degrees of male-female expression balance, and potentially also in their corresponding degrees of dosage compensation in males. Individual sex-linked genes can vary in whether they exhibit Type I-IV expression profiles (Saunders and Muyle 2024; Webster et al. 2024), even in squamate reptiles with mechanisms for near-complete dosage compensation, such as *Anolis carolinensis* (Marin et al. 2017; Rupp et al. 2017). Interestingly, the same Type IV pattern of male-female expression imbalance appears to characterize the more recently X-linked distal region of the *S. undulatus* X chromosome (DR10). However, it is unclear whether Y degeneration has proceeded to the extent that males actually require dosage compensation for most of this neo-X region, given that log_2_ male/female DNA coverage ranges from 0 to -1 for individual genes in DR10 and that not at all genes may actually require compensation given their biological function.

In contrast to the ancestral region of the X chromosome, the distal region of *S. undulatus* chromosome 6 (DR6) that we identify as a putative neo-X region in *S. jarrovii* appears to exhibit an overall “Type III” expression profile, corresponding to the lack of dosage compensation in males and the lack of expression balance between sexes. This pattern is common in ZW systems, including other squamate reptiles (Rovatsos et al. 2019; Webster et al. 2024), but rare in XY systems (Gu and Walters 2017; Saunders and Muyle 2024). Nonetheless, our comparisons of DR6 expression against randomly sampled autosomal blocks from *S. jarrovii* and also against orthologs in the ancestral autosomal state in *S. undulatus* both indicate that DR6 in *S. jarrovii* is expressed at typical autosomal levels in females and at substantially lower levels in males. Such Type III expression profiles may correspond to early evolutionary stages in which degeneration of Y has created dosage effects in males, but mechanisms for compensatory upregulation of X have yet to evolve. Likewise, Type IV expression profiles may correspond to later evolutionary stages in which dosage compensation has been achieved via the upregulation of X in males, but male-female expression imbalance persists due to the concomitant upregulation of X in females. Ohno (1967) hypothesized that this state may precede the evolution of mechanisms such as X-inactivation in females, which then restore male-female expression balance (Grant et al. 2012; Marin et al. 2017). However, it is unclear whether “Type I” patterns characterized by dosage compensation and male-female expression balance actually evolve via this proposed sequence, particularly in vertebrates (Gu and Walters 2017; Saunders and Muyle 2024). While many ancestrally X-linked genes do exhibit both dosage compensation and male-female expression balance in *S. undulatus*, it is clear that many others still lack male-female expression balance despite sufficient time for its evolution in this ancient (>100 Mya) XY system.

The anolid lizard *Anolis carolinensis* has a *Drosophila*-like epigenetic mechanism for the male-specific upregulation of hemizygous X-linked genes (Marin et al. 2017), producing a Type I pattern of both dosage compensation and male-female expression balance across the majority of the X chromosome (Chen et al. 2020a; Rovatsos and Kratochvíl 2021; Rupp et al. 2017). This is noteworthy as one of the few examples of near-complete dosage compensation accompanied by near-complete male-female expression balance in any vertebrate (Chen et al. 2020a; Metzger et al. 2021; Saunders and Muyle 2024). Intriguingly, recent analyses of the phrynosomatid lizards *Urosaurus nigricaudus* and *U. bicarinatus* suggested both dosage compensation and male-female expression balance across X, raising the possibility that the epigenetic mechanism for male-specifc upregulation of X that was first discovered in *A. carolinensis* may be broadly conserved across these distantly related families in Pleurodonta that share a homologous X chromosome (Davalos-Dehullu et al. 2023). Our results are only partly consistent with this interpretation. Although we find that the ancestral region of X does exhibit complete (brain, muscle) to partial (liver) dosage compensation relative to autosomes in *S. undulatus* males, we also find clear evidence of widespread male-female expression imbalance for this same ancestral region of X across phrynosomatids, including another *Urosaurus* species, *U*. *ornatus*.

This discrepancy between studies is unlikely to reflect differences in the genomes used for read mapping (*U. nigricaudus* versus *S. undulatus*), as both produce clear signatures of male hemizygosity for the ancestral region of X when mapping genomic DNA. However, whereas we set out to explicitly test for differential expression on a gene-by-gene basis by calculating mean male and female expression levels from multiple replicate libraries (*n* = 5-7 per sex) across multiple ages and tissues, Davalos-Dehullu et al. (2023) derived male-female expression ratios from pairwise combinations of one individual male library against one individual female library (*n* = 2 libraries total per sex). This difference in methodology and sample size, potentially in combination with our use of isolated tissues (brain, liver, and muscle) rather than tail samples comprised of multiple tissue types (*e.g.,* bone, muscle, skin), may have afforded us greater power and precision for identifying sex-biased expression. Importantly, we note that the majority of genes on X lack sex-biased expression for most ages and most tissues, and several even exhibit consistently male-biased expression despite their hemizygosity in males. Therefore, much of the X chromosome does appear to exhibit both dosage compensation and at least partial male-female expression balance in phrynosomatids, despite the clear lack of male-female expression balance for dozens of individual genes. Even those genes that are consistently female-biased across ages and tissues have mean expression ratios of 1.7 rather than 2-fold, indicating partial expression balance. In this sense, *S. undulatus* and other phrynosomatids appear similar to many other species in which dosage compensation is partial and occurs on a gene-by-gene basis (Furman et al. 2020; Gu and Walters 2017; Mank 2013, 2009; Mank and Ellegren 2009; Mullon et al. 2015). This is true even for the near-complete dosage compensation and male-female expression balance seen in *A. carolinensis* (Marin et al. 2017; Rupp et al. 2017; Saunders and Muyle 2024).

Chromosomal fusions are common in *Sceloporus* (Leaché et al. 2016), particularly for sex chromosomes (Lisachov et al. 2021). It has been proposed that pseudoautosomal regions (PARs) on the tips of sex chromosomes may facilitate such fusions by avoiding the creation of multiple sex chromosome systems, since the newly fused, formerly autosomal segment can be transferred to the other sex chromosome (*e.g*., from X to Y) by recombination (Lisachov et al. 2021). Although we cannot assess the structure of the putative X-autosome translocation from chromosome 6 to 10 in *S. jarrovii*, the fusion that formed DR10 in the *S. undulatus* lineage is located adjacent to a small region of ancestrally X-linked genes with balanced DNA coverage and expression between the sexes, suggesting that the fusion occurred at a terminal PAR. Interestingly, many of the same syntenic blocks of genes have been repeatedly involved in fusions with sex chromosomes across squamate reptiles (Lisachov et al. 2021), and the DR10 region in *S. undulatus* provides another example of this phenomenon. DR10 maps to a portion of a putatively autosomal michrochromosome in other phrynosomatid genomes, but the entire microchromosome has undergone an independent fusion with X in the anolid *A. sagrei* (Geneva et al. 2022; Giovannotti et al. 2017; Kichigin et al. 2016).

Finally, our study illustrates how the transcriptomic signatures of male-female expression imbalance can both generate and test hypotheses about the evolution of sex chromosomes, even in the absence of genomic or karyotypic data. For example, the phylogenetically conserved pattern of male-female expression imbalance for genes that map to the ancestrally X-linked region of *S. undulatus* chromosome 10 supports the hypothesis that this region has remained largely X-linked throughout phrynosomatid evolution, with the possible exception of *S. sleveni*.

By contrast, the distal region of chromosome 10 (DR10) only exhibits this signature of male-female expression imbalance in *S. undulatus,* consistent with the hypothesis that this region became X-linked more recently, after the split between lineages that gave rise to *S. undulatus* and *S. virgatus*. Syntenty analyses indicate that this region is also X-linked in *S. tristichus*, suggesting that it fused with the ancestral X chromosome approximately 6-10 Mya based on divergence estimates for these three species (Leaché et al. 2016; Westfall et al. 2021). We also found a strong signature of male-female expression imbalance in *S. jarrovii* for genes that map to a small region of *S. undulatus* chromosome 6 (DR6), and the absence of this signature from all other species in our analysis supports the hypothesis that DR6 is also a recently X-linked region. Given that male-female expression imbalance is pronounced and male expression appears to lack dosage compensation for DR6 in *S. jarrovii*, this may represent a younger neo-X region than DR10. However, our current transcriptome sampling across *Sceloporus* lacks the phylogenetic resolution to estimate its age more precisely. In future comparative work, tests for transcriptomic signatures of male-female expression imbalance could provide an efficient way to identify and date similar X-autosome fusions in this and other lineages, particularly when sequencing tissues or ages that exhibit relatively little sex-biased expression from other regulatory mechanisms.

## Materials and Methods

### Synteny and Dosage of the *S. undulatus* X Chromosome

To confirm inferences from several previous studies (Davalos-Dehullu et al. 2023; Westfall et al. 2021), we assessed the synteny of *S. undulatus* chromosome 10 (NC_056531.1) with the genomes of three other iguanian lizards in the Pleurodonta clade that share an ancient (>100 Mya) X chromosome: *Phrynosoma platyrhinos* (GCA_020142125.1, MUOH_PhPlat_1.1, (Koochekian et al. 2022)), *Anolis carolinensis* (GCF_000090745.2, AnoCar2.0, (Alföldi et al. 2011)), and *Anolis sagrei* (GCA_025583915.1, AnoSag2.1, (Geneva et al. 2022)). Each of these other three genomes was sequenced and assembled from a female specimen. The ancestral X chromosome for this lineage has undergone multiple fusions with autosomes to form several more recently X-linked strata in *A. sagrei* (Geneva et al. 2022). Given the phylogenetic distance separating the species in our analyses (common ancestor ca. 83 Mya), we determined orthology-based syntenic relationships using the R package *GENESPACE* (Lovell et al. 2022), which aligns amino acid sequences using *DIAMOND2* (Buchfink et al. 2015), identifies orthogroups with *OrthoFinder* (Emms and Kelly 2019, 2015), and determines syntenic regions for each non-reciprocal pair of genomes by collapsing syntenic orthogroups into a pan-genome annotation.

*GENESPACE* identified a minimum of 16,657 genes distributed across 15,662 orthogoups in *P. platyrhinos* and a maximum of 28,605 genes distributed across 15,458 orthogroups in *A. carolinensis* after filtering for overdispersion. We used the *GENESPACE* function ‘plot_riparian’ to visualize synteny for the *S. undulatus* genome with specific focus on chromosome 10.

To precisely resolve synteny of the X chromosome at the same gene-by-gene resolution as our transcriptomic analyses, we conducted BLASTN (Altschul et al. 1990; Madden 2013) searches using the longest annotated transcript for each gene on *S. undulatus* chromosome 10 (*n* = 330 with XM accessions) as a query against the NCBI reference genomes of five other iguanian lizards in Pleurodonta: closely related *Sceloporus tristichus,* which was formerly considered a subspecies of *S. undulatus* (GCA_016801065.1, ASM1680106v1 (Bedoya and Leaché 2021)), the additional phrynosomatids *Urosaurus nigricaudus* (GCA_034509955.1, ASU_Unig_1 (Davalos-Dehullu et al. 2023)) and *Phrynosoma platyrhinos* (GCA_020142125.1, MUOH_PhPlat_1.1 (Koochekian et al. 2022)), and two more distantly related anolids, *Anolis sagrei* (GCA_037176765.1, rAnoSag1) and *A. carolinensis* (GCA_035594765.1, rAnoCar3.1). We used default BLASTN parameters with minor modifications (reward = 1, penalty = 2, and word_size = 28) and sorted results by lowest e-value and highest query coverage. Across every search, the top three hits were always from the same gene, so we used the top BLASTN hit for each transcript to infer the chromosomal location of the homologous gene in each of the other five species.

To characterize sex differences in dosage, we used NCBI *sratoolkit* v.3.0.5 to download short-read, whole-genome sequences (Illumina paired-end 150 bp reads) from 18 female and 11 male *S. undulatus* specimens obtained by Assis et al. (2025) as part of a separate study (NCBI accession numbers SRR12504882 to SRR12504910 within bioproject PRJNA656311). We trimmed adaptors and filtered low-quality reads used *fastp* (Chen et al. 2018) with default settings, then summarized quality control metrics using *multiqc* (Ewels et al. 2016). Samples had a mean duplication rate of 8.8% and we retained a mean of 319 M reads per sample (>95% of input) after filtering. We aligned reads to the *S. undulatus* genome using *bowtie2* (Langmead and Salzberg 2012) with default settings and removed alignments with a mapping quality (MQ) score below 20 using *samtools* (Li et al. 2009). The alignment rate exceeded 95% for all specimens.

We estimated per-base pair depth of coverage for each individual with ‘genomeCoverageBed’ and calculated the mean depth of coverage for each sex over non-overlapping 100-kbp windows across the entire genome using ‘map’ from *bedtools* (Quinlan and Hall 2010). To correct for differences in sequencing depth across individuals, we weighted each 100-kbp window value by the genome-wide mean depth of coverage for that individual using custom *awk* functions. Individuals were sequenced at 54x coverage on average, with normalized depth of coverage ranging from 0.88 (chromosome 11) to 1.02 (chromosome 8) in females and 0.56 (chromosome 10) to 1.03 (chromosome 8) in males.

To calculate a per-gene weighted coverage value, we first divided the mean coverage depth across the open reading frame of each gene by the mean depth of coverage of its chromosome or scaffold in each individual. Next, we caculated the mean weighted coverage per gene, per sex. We then calculated log_2_ (mean weighted coverage in males + 0.001 / mean weighted coverage in females + 0.001) for each gene, which we refer to as log_2_ (male/female) coverage. X-linked genes that are hemizygous in males should have log_2_ (male/female) coverage values near -1. Although the *S. undulatus* genome was sequenced from a male specimen, it does not contain an assembled Y chromosome (Westfall et al. 2021). To identify potential Y-liked genes, we searched for any scaffolds containing at least one annotated gene with a mean log_2_ (male/female) coverage value greater than 0.5. To identify potential X-liked genes that were not assembled with chromosome 10, we searched for any scaffolds containing at least one annotated gene with a mean log_2_ (male/female) coverage value less than -0.5.

### Tissue Collection for Gene Expression

To describe sex differences in gene expression, we collected tissues immediately upon capture of wild animals in their natural habitats. All procedures were reviewed and approved by the Institutional Animal Care and Use Committees at the University of Virginia (protocol 3896), Florida International University (IACUC-20-012-CR01), and Rutgers University (PROTO 201800157). Animals were collected under permits from Arizona Game and Fish (SP658032 and SP407104), Texas Parks and Wildlife (SPR-0719-113), and New Jersey Fish and Wildlife (SC 2019007). Additional collecting and land use permissions were obtained from Catalina State Park (AZ), the Coronado National Forest branch of the United States Forest Service (AZ), Tucson Parks and Recreation (AZ), Dalquest Desert Research Station (TX), and Colliers Mills Wildlife Management Area (NJ). Collecting locations are provided in Table S1.

We collected *S. undulatus* and *S. jarrovii* at each of three ages: neonate, maturing, and adult (*n* = 5-7 individuals per age, per sex; Table S1). Neonates were collected within several weeks of hatching in oviparous, spring-breeding *S. undulatus* (early September, age <1 mo) and within several weeks of birth in viviparous, fall-breeding *S. jarrovii* (mid June, age <1 mo). Maturing individuals were sampled when sex differences in growth, coloration, and circulating testosterone first emerge in the summer after hatching in *S. undulatus* (July, age 10-11 mo) and the first fall breeding season after birth in *S. jarrovii* (late September, age 4-5 mo) (Cox and John-Alder 2007, 2005; Cox et al. 2005). Reproductive adults were sampled during the breeding season in both *S. undulatus* (mid May, age >20 mo) and *S. jarrovii* (late September, age >16 mo). To assess the evolution of sex-biased gene expression, we also collected reproductive adults of five additional *Sceloporus* species (*S. clarkii, S. magister, S. merriami, S. slevini, S. virgatus*) and three species representing other phrynosomatid genera (*Cophosaurus texanus, Urosaurus ornatus*, *Uta stansburiana*; *n* = 6 per sex, per species). Adults of each species were sampled in late April or early May during their breeding seasons (Table S1).

We captured lizards by hand or using a hand-held pole with a loop of braided fishing line, then immediately measured them for mass (nearest 0.1 g) and snout-vent length (nearest 1 mm). Sex was determined according to presence or absence of enlarged post-anal scales (present only in males) and confirmed by inspection of the gonads during tissue collection. Immediately following measurement, we euthanized lizards via decapitation and collected liver, whole brain, and muscle (along the femur) into RNAlater (Qiagen, Valencia, CA, USA) on ice. Tissues were held overnight at 4°C, frozen at -20°C, and then stored at -80°C until RNA extraction.

### RNA Extraction and Sequencing

We extracted total RNA from liver for all 10 species (Table S1). For *S. undulatus* and *S. jarrovii*, this included three age groups (neonate, maturing, and adult). For *S. undulatus*, we also extracted RNA from brain and muscle from all three age groups. RNA was extracted using the Illustra/Cytiva RNAspin Mini RNA Isolation Kit (Global Life Sciences Solutions USA, Marlborough, MA, USA), following the manufacturer’s protocol, with the exception that all tissues were homogenized twice (45 s per round at 30 Hz) in lysis buffer using 5-mm stainless steel beads in a Tissuelyser II bead mill (Qiagen; Germantown, MD, USA). We assessed RNA concentrations for all samples using a Nanodrop spectrophotometer (ThermoFisher Scientific, Waltham MA, USA). We also assessed RNA quality and concentration for most samples on an Agilent Bioanalyzer (RNA 6000 Pico; Agilent Biotechnologies, Santa Clara, CA, USA). All samples exhibited sufficient quality for library preparation and sequencing (mean RIN = 8.1).

Preparation of cDNA libraries and RNA sequencing was carried out by the Georgia Genomics and Bioinformatics Core at the University of Georgia (Athens, GA, USA). Libraries were prepared using the KAPA Stranded mRNA-Seq kit for Illumina platforms with poly(A) selection (KAPA Biosystems; Wilmington, MA, USA). Library preparation and sequencing were conducted in three batches with respect to species and tissue (Table S1) to optimize comparisons between sexes and to obtain approximately 20 M reads per sample. In the first batch, libraries for *S. undulatus* and *S. jarrovii* liver from all three ages were prepared together using ∼1 μg total RNA per sample. In the second batch, libraries for *S. undulatus* brain and muscle from all three ages were prepared using ∼2 μg total RNA (brain) or ∼0.5 μg total RNA (muscle), with libraries from each tissue loaded in equimolar ratios for sequencing. In the third batch, libraries for *S. virgatus, S. clarkii, S. slevini, S. magister, S. merriami*, *U. ornatus*, *U. stansburiana*, and *C. texanus* liver from adults were prepared using ∼1 μg total RNA. Each batch was sequenced twice on Illumina P3 flow cells (paired-end, 100-bp reads) using an Illumina NextSeq 2000.

### Read Count Generation and Differential Expression Analysis

All RNAseq reads were assessed for quality and trimmed using *fastp* (Chen et al. 2018) with paired-end base correction, low-complexity filtering, and 3’ end tail quality trimming (average phred score threshold = 20 and window size = 5). Poly(g) and poly(x) trimming were also enabled to remove low complexity sequences from the 3’ end of reads, along with an overall minimum length filter of 36 bp and minimum phred quality threshold of 25. Cleaned read pairs and reads orphaned during trimming (*i.e*., read partner removed due to length or quality) were aligned to the *S. undulatus* genome (GCA_019175285.1; SceUnd_v1.1) using *subread-align* (Liao et al. 2013b) with *S. undulatus* RefSeq coding sequences (GCF_019175285.1) as a guide during alignment (accessed via NCBI). Although we used the *S. undulatus* genome as a common reference to generate comparable read counts across all 10 species, we note that the genus *Sceloporus* exhibits widespread chromosomal evolution (Leaché et al. 2016), such that gene locations in other species may differ from those in the *S. undulatus* assembly.

For all reads from *Sceloporus* species, we used default parameters in *subread-align* to identify valid alignments to coding regions in the *S. undulatus* genome. For *Urosaurus*, *Uta*, and *Cophosaurus*, we relaxed alignment stringencies due to increased phylogenetic distance and reduced alignment rates observed when default parameters were used. This entailed a lower minimum number of subreads necessary to report an alignment (-m 2), an increased number of mismatches allowed (-M 5), a lower minimum fragment size (-d 35), an increased maximum fragment size (-D 750), and an increased maximum number of indels (-I 7). Paired reads and reads orphaned during *fastp* trimming were aligned independently and expression counts for all species were generated from alignments using *featureCounts* (Liao et al. 2013a) with default parameters. We summed expression counts from paired reads and orphaned reads to generate final expression estimates for each sample. Only unique alignments were counted towards the total expression estimate for a gene (*i.e.,* multi-mapping reads were not counted).

We conducted differential expression analyses using R (v 4.5.2) (R Core Team, 2019) package *edgeR* (v 4.8.2) (Chen et al. 2016; McCarthy et al. 2012; Robinson et al. 2009; Robinson and Smyth 2007). Prior to analysis, we filtered counts to remove low-expression genes using *edgeR* function ‘FiltrByExpr’ separately for each tissue within each species (Table S1, “edgeR model”). To identify and remove outlier libraries after low-expression filtering, we applied a TMM (trimmed mean of M-values) normalization to expression counts in *edgeR*, then used a log_10_ counts-per-million (CPM) transformation to adjust for library size. We used robust PCA on log_10_ (CPM) values to identify and remove any outlier libraries using R package *rrcov* and function ‘PcaGrid’ (crit.pca.distance = 0.975) (Chen et al. 2020b). We conducted outlier analyses separately for each tissue within each species. The total numbers of samples passing outlier filtering are reported in Table S1. Following outlier removal, we conducted differential expression analysis using TMM-normalized read counts, which we fit to a negative binomial model (*edgeR* function ‘glmQLFit’, robust = TRUE) with separate models for each species. We identified sex-biased DEGs (differentially expressed genes) using quasi-likelihood *F-*tests comparing female versus male expression for each species with *edgeR* function ‘glmQLFTest’. Within each species, we made pairwise comparisons between sexes for each combination of age and tissue. Genes with a Benjamini-Hochberg *P*-value adjusted for the transcriptome-wide false discovery rate (“FDR”) < 0.05 were considered DEGs (Benjamini and Hochberg 1995).

### Testing for Male-Female Expression Imbalance in *S. undulatus*

After identifying significantly sex-biased DEGs within each combination of age and tisse, we identified the subsets of genes that are consistently female-biased or consistently male-biased DEGs across all ages within a tissue, then determined the chromosomal locations of these genes in the *S. undulatus* assembly. If consistently female-biased DEGs are the result of male-female dosage imbalance, then these DEGs should map predominantly to chromosome 10, the X chromosome. If consistently male-biased DEGs are the result of Y-linkage, then these DEGs should map predominantly to unplaced scaffolds in the *S. undulatus* assembly, which does not include an assembled Y chromosome but does contain sequence from a male specimen. Mapping of consistently sex-biased DEGs to autosomes (chromosomes 1-9 and 11) would indicate either an error in sex chromosome assembly, or a mechanism for consistently sex-biased expression other than dosage imbalance due to male hemizygosity. As noted above, we assessed log_2_ male/female DNA coverage for each of these consistently sex-biased DEGs to infer underlying patterns of dosage. We also visually assessed genome-wide patterns of sex-biased expression using Mahattan plots with the -log_10_ (*P-*value) of differential expression for each gene ordered sequentially by chromosome number and transcriptional start site.

As a complementary approach to our DEG analysis, we assessed chromosome-wide patterns of sex-biased expression by using *edgeR* function ‘rpkm’ to calculate the expression of each gene as FPKM (fragments per kilobase of transcript per million mapped reads) using *S. undulatus* gene lengths (GCF_019175285.1). For each tissue, we used Wilcoxon signed-rank tests with a Bonferroni-adjusted alpha (0.05 / 3 = 0.017) to test whether the median sex bias in expression (log_2_ male/female FPKM) of genes on chromosome 10 differed significantly from 0. We conducted these analyses both with and without the subsets of consistently sex-biased DEGs previously identified in each tissue (see above). Because the current *S. undulatus* assembly does not include a unique Y scaffold, we could not conduct similar tests for Y-linked expression.

### Reassesssing Male-Female Expression Imbalance in *S. undulatus* Using the *S. tristichus* Genome

Because the assembly of the *S. undulatus* X chromosome may have been influenced by homologous reads from Y (Westfall et al. 2021), we repeated our analyses of differential gene expression by aligning RNAseq reads from *S. undulatus* to the genome of a closely related congener (*Sceloporus tristichus,* formerly *S. undulatus tristichus*) that was sequenced exclusively from female tissue (Bedoya and Leaché 2021). Using this assembly should minimize any artifacts potentially introduced by similar X and Y sequences in the *S. undulatus* genome, such as the co-assembly of X and Y gametologs on chromosome 10 or the failure of X gametologs to assemble on chromosome 10. Read alignments and estimates of gene expression were made using subread-align and featureCounts against the *S. tristichus* genome (ASM1680141v1) and gene annotations from Bedoya and Leaché (2021; Sceloporus_tristichus_snow_11.gff3) with the same parameters used during alignment to the *S. undulatus* genome (described above). Results of these analyses are presented in the Supplementary Materials.

### Identification of Potential Gametolog Pairs in *S. undulatus*

If males express two formerly allelic copies of the same gene (*i.e*., X and Y gametologs), then the independent mapping of reads to each gametolog could create the appearance of female-biased expression of the X gametolog for a gene that is otherwise expressed similarly in both sexes. For all genes identified as consistently sex-biased DEGs using the *S. undulatus* genome, we used BLASTN (Madden 2013) to search for similar sequences that could represent potential gametologs. We used the longest annotated transcripts for each consistently female-biased DEG as query sequences and searched them against the *S. undulatus* (taxid:8520) refseq_rna database using default alignment parameters (E-threshold = 0.05; word size = 11; max matches in query = 0; match/mismatch scores = 2,-3; gap costs = Existence: 5, Extension: 2) with low-complexity filters enabled. We queried the top 100 hits for each female-biased DEG sequence to identify any located on unassembled scaffolds, then assessed male and female expression levels (FPKM) to identify those with low expression in females, indicating possible Y gametologs. We applied a similar strategy to identify any potential X gametologs for consistently male-biased DEGs, which would be expected to occur on chromosome 10 or unplaced scaffolds. In situations where we identified a potential gametolog pair, we re-assessed patterns of sex bias after pooling reads from each separately assembled region (*i.e*., summing putative X and Y gametolog reads). Results of these analyses are presented in the Supplementary Materials.

### Comparing X-linked to Autosomal Expression in *S. undulatus*

We established null distributions of median autosomal expression by randomly sampling spatially contiguous blocks of the same number of autosomal genes as expressed genes on X. We calculated the median log_2_ FPKM expression for each block of genes in each tissue, age, and sex, repeating this random selection process 1000 times throughout the autosomal genome to generate a null distribution of median autosomal expression for each combination of tissue, age, and sex. For each sampling iteration, we selected a contiguous block of genes on the same scaffold (excluding genes on unplaced scaffolds) and arrayed consecutively by start position, similar to the arrangement of genes on X. Each block of autosomal genes was sampled only once to avoid pseudoreplication. We then tested whether the median log_2_ FPKM expression of genes on X fell outside the lower 5% or upper 95% quantiles of the corresponding autosomal distribution. If median expression on X falls below the corresponding null distribution of median autosomal expression in males, this would indicate a lack of dosage compensation (“Type III” expression pattern, Gu and Walters 2017). If the median expression of genes on X falls above the corresponding null distribution in females, this could indicate that dosage compensation in males has resulted in overexpression of X in females (“Type IV” expression pattern; Gu and Walters 2017). We conducted these tests separately for genes on the ancestrally X-linked, 14.85-Mbp region of chromsosome 10 (*n* = 231 to 255 expressed genes, depending upon tissue), as well as the more recently X-linked, 1.76-Mbp distal region of chromosome 10 (*n* = 41 to 46 expressed genes), with null distributions of autosomal expression generated separately for each region based on its number of expressed genes in each tissue.

### Testing for Male-Female Expression Imbalance in *S. jarrovii*

For *S. jarrovii*, we used the same methods described above for *S. undulatus* to identify consistently sex-biased genes in the liver (i.e., significant DEGs biased towards the same sex across all three ages), and to test for overall male-female expression imbalance across genes on chromosome 10 in the *S. undulatus* assembly (*i.e*., significant deviations of median log_2_ male/female FPKM ratios from 0 using Wilcoxon signed-rank tests with Bonferroni correction for tests across 3 ages). Because the distal 1.76-Mbp portion of *S. undulatus* chromosome 10 (DR10) is presumably autosomal in *S. jarrovii*, we tested for male-female expression imbalance separately in DR10 and in the ancestrally X-linked 14.85-Mbp region of chromosome 10, which we expected to be X-linked in both species.

Our analyses in *S. jarrovii* unexpectedly uncovered a signal of highly sex-biased expression in genes mapping to a small (∼11 Mbp) distal region on chromosome 6 in the *S. undulatus* assembly (hereafter “DR6”), so we conducted additional analyses to test whether this region exhibits the transcriptomic signatures of an X chromosome in *S. jarrovii*. We used Wilcoxon signed-rank tests (as described above for chromosome 10) to test whether the median log_2_ male/female FPKM expression ratio for genes on DR6 differs from 0, and also to compare (separately within each sex) the median log_2_ FPKM expression of genes in DR6 to null distributions of median log_2_ FPKM expression for 1000 randomly sampled genes located contiguously in other autosomal regions. For comparison, we conducted similar analyses of DR6 in *S. undulatus,* where it is autosomal. To directly test for an evolutionary reduction in expression of DR6 genes in *S. jarrovii* males, as expected if this formerly autosomal region has become X-linked but still lacks complete dosage compensation, we conducted paired (by gene) *t*-tests comparing expression of DR6 homologs between *S. undulatus* and *S. jarrovii*. We predicted that expression of DR6 would be similar in females of each species, but reduced in *S. jarrovii* males relatieve to *S. undulatus* males.

### Characterizing the Evolution of Gene Expression Across Phrynosomatidae

We generated liver transcriptomes for adult males and females of 7 *Sceloporus* species, plus representatives of 3 other phrynosomatid genera (*n* = 5-7 individuals per sex, per species; Table S1). For species, we tested whether genes that map to the ancestrally X-linked, male-hemizygous, 14.85-Mbp region of chromosome 10 in the *S. undulatus* assembly exhibit female-biased expression. To resolve the evolution of neo-X regions, we also tested whether genes that map to the distal regions of *S. undulatus* chromosome 10 (DR10, putative neo-X region in *S. undulatus*) and chromosome 6 (DR6, putative neo-X region in *S. jarrovii*) exhibit female-biased expression in any other species. For each comparison described above, we used a 1-sample Wilcoxon signed-rank test with Bonferroni correction for the 10 comparisons across species (adjusted alpha = 0.005) to test for male-female dosage imbalance (median log_2_ male/female FPKM < 0) of all expressed genes that map to the corresponding regions of the *S. undulatus* assembly. We also conducted similar analyses using only the subsets of consistently female-biased DEGs from chromosome 10 (*i.e.,* DEGs in *S. undulatus* transcriptomes) or DR6 (*i.e.,* DEGs in *S. jarrovii* transcriptomes). We visualized evolutionary patterns in gene expression using a consensus phylogeny based on several sources (Leaché et al. 2016; Pyron et al. 2013).

## Supporting information

Hale et al. Supplemental Material

## Acknowledgments

We thank Matt Watson and Tyler Wittman for help with sample collection. We thank the Georgia Genomics and Bioinformatics Core (GGBC) for library preparation and sequencing. The authors acknowledge Research Computing at The University of Virginia (https://rc.virginia.edu) for providing compulational resources and technical support that have contributed to the results reported within this publication. This work was supported by a Collaborative Research award from the U.S. National Science Foundation (IOS 1755134 to CLC, IOS 1754934 to HBJ-A, and IOS 1755026 to RMC). CDR was supported by a Research Traineeship from the U.S. National Sceince Foundation (NRT-ROL 2021791). PLHM was supported by a Rising Scholars Postdoctoral Fellowship from the University of Virginia.

## Data Availability

RNAseq data underlying this article will be made available through the National Center for Biotechnology Information (NCBI) Short Read Archive upon acceptance. Read counts and code, as well as edgeR DEG output and FPKM expression estimates from edgeR, are available via figshare (https://doi.org/10.6084/m9.figshare.30324850).

