## Supplementary material for "Widespread male-female expression imbalance of X-linked genes across phrynosomatid lizards": Hale et al. Supplemental Material

### Supplementary Materials for: Widespread male-female expression imbalance of X-linked genes across phrynosomatid lizards

Matthew D. Hale[1,2,*], Pietro H. de Mello[1], Daniel T. Nondorf[1], Christopher D. Robinson[1,3], Henry B. John-Alder[2], Christian L. Cox[3], and Robert M. Cox[1,*]

[1] Department of Biology, University of Virginia, Charlottesville, VA 22904

[2] U.S. Military HIV Research Program, Walter Reed Army Institute of Research, Silver Spring, MD 20910; Henry M. Jackson Foundation for the Advancement of Military Medicine, Inc., Bethesda, MD, 20817

[3] School of Life Sciences, Arizona State University, Tempe, AZ 85287

[4] Department of Ecology, Evolution, and Natural Resources, Rutgers University, New Brunswick, NJ 08901

[5] Department of Biological Sciences, Florida International University, Miami, FL 33199

**Authors for Correspondence:**

*Matthew D. Hale: Washington, DC 20009, USA.

*Robert M. Cox: Department of Biology, University of Virginia, Charlottesville, VA 22904, USA. Phone: 434-982-1987.

#### Supplementary Results

<u>Confirming Male-Female Expression Imbalance in *S. undulatus* Using the *S. tristichus* Genome</u>

The genome-wide patterns of sex-biased expression that we described in *S. undulatus*, including inferred male-female expression imbalances on X, were recapitulated when we repeated our analyses by mapping *S. undulatus* reads to the genome of *S. tristichus* (formerly *S. undulatus tristichus*), a closely related congener with a genome sequenced exclusively from female tissue (Bedoya and Leaché, 2021). Across liver, brain, and muscle, consistently female-biased DEGs ($n$ = 42) exhibit a relatively narrow range of $\log_2$ FC values averaging 1.7-fold higher expression in females (Fig. S7) and localize exclusively to chromosome 10, the predicted *S. tristichus* X chromosome (Fig. S8). Moreover, median expression of all genes on X is significantly female-biased across all ages and tissues when mapped to the *S. tristichus* assembly (Fig. S8). The few consistently male-biased DEGs that we detected ($n$ = 19) exhibit a broad range of extreme $\log_2$ FC values and localize to both X and autosomes (Fig. S8). Genes on unassembled scaffolds are not annotated in the *S. tristichus* assembly, which could explain why we detected slightly fewer consistently female-biased DEGs relative to our analyses using the *S. undulatus* assembly. This plus the absence of Y-linked sequence from the *S. tristichus* assembly likely explains why we detected substantially fewer consistently male-biased DEGs. Although it is unclear why several consistently male-biased DEGs in *S. undulatus* brain map to autosomes in the *S. tristichus* assembly (Fig. S8), the fact that all of the significantly female-biased DEGs that we identified map to chromosome 10 in the *S. tristichus* assembly lends additional support (along with ratios of DNA coverage indicating that males are hemizygous for most ancestrally X-linked genes outside of a small pseudoautosomal region; Fig. 1) to the inference that the enrichment of sex-biased expression on X is not driven by any artifacts arising from the co-assembly of X and Y sequences in the *S. undulatus* assembly.

Identification of Potential Gametolog Pairs in *S. undulatus*

We tested for potential X and Y gametologs by conducting BLAST searches of each consistently sex-biased DEG against the entire *S. undulatus* refseq-RNA database. We found significant hits that map to unassembled scaffolds for 10 of 45 consistently female-biased genes and 2 of 10 consistently male-biased genes on the X chromosome (12 of 55 total), and for 5 of 15 consistently female-biased genes and 20 of 31 consistently male-biased genes on unplaced scaffolds (25 of 46 total). However, none of the hits for consistently female-biased genes on X exhibit highly male-biased expression, as would be expected for Y gametologs, and neither of the hits for consistently male-biased genes on X exhibit the reciprocally expected pattern of female-biased expression, as they are not consistently female-biased across all 3 ages and also exhibit very little sequence overlap (~50 bp) with the query DEG sequence. Despite the large number of BLAST hits for sex-biased genes on unplaced scaffolds, we only identified three gene pairs for which these hits correspond to a gene with significant sex-biased expression opposite that of the query sequence (Fig. S9). In each of these three pairs, both genes are located on unplaced scaffolds and may represent gametologs, although a low (non-zero) level of expression in females is observed in the putative Y gametolog for two of these pairs (Fig. S9). Aside from these three gene pairs, most BLAST hits we identified appear to correspond to unassembled autosomal loci with relatively low sequence overlap between the DEG and its BLAST hits, rather than orphaned gametolog pairs in the *S. undulatus* assembly.

**Table S1.** Summary of samples collected and sequenced in this study. "RNAseq batch" indicates libraries prepared and sequenced together on the same flow cell. "EdgeR model" indicates filtered read counts grouped together for low-expression gene filtering, outlier detection, and model construction in edgeR. "*N* sampled (retained)" indicates the number of individuals collected and the number of libraries retained for analysis after exclusion of outliers.

| Species | Site | Tissue | Age | Collection date | RNAseq batch | EdgeR model | *N* sampled (retained) | |
|---|---|---|---|---|---|---|---|---|
| | | | | | | | Female | Male |
| *S. undulatus* | NJ[1] | Liver | Neonate | Sep 2019 | 1 | 1 | 6 (6) | 6 (6) |
| | | | Maturing | Jul 2019 | 1 | 1 | 6 (6) | 6 (6) |
| | | | Adult | May 2019 | 1 | 1 | 6 (5) | 7 (7) |
| | | Brain | Neonate | Sep 2019 | 2 | 2 | 6 (6) | 6 (6) |
| | | | Maturing | Jul 2019 | 2 | 2 | 6 (6) | 6 (5) |
| | | | Adult | May 2019 | 2 | 2 | 6 (5) | 7 (7) |
| | | Muscle | Neonate | Sep 2019 | 2 | 3 | 6 (4) | 6 (6) |
| | | | Maturing | Jul 2019 | 2 | 3 | 6 (5) | 6 (5) |
| | | | Adult | May 2019 | 2 | 3 | 6 (6) | 7 (6) |
| *S. jarrovii* | AZ[2,3] | Liver | Neonate | Jun 2019 | 1 | 4 | 6 (5) | 6 (5) |
| | | | Maturing | Sep 2019 | 1 | 4 | 6 (4) | 6 (6) |
| | | | Adult | Sep 2019 | 1 | 4 | 5 (5) | 5 (5) |
| *S. clarkii* | AZ[4] | Liver | Adult | May 2021 | 3 | 5 | 6 (6) | 6 (6) |
| *S. magister* | AZ[4,5] | Liver | Adult | May 2021 | 3 | 6 | 6 (6) | 6 (6) |
| *S. merriami* | TX[6] | Liver | Adult | Apr 2021 | 3 | 7 | 6 (6) | 6 (6) |
| *S. slevini* | AZ[7] | Liver | Adult | May 2021 | 3 | 8 | 6 (6) | 6 (6) |
| *S. virgatus* | AZ[2] | Liver | Adult | May 2021 | 3 | 9 | 6 (6) | 6 (5) |
| *U. ornatus* | AZ[8] | Liver | Adult | May 2021 | 3 | 10 | 6 (6) | 6 (6) |
| *U. stansburiana* | AZ[9] | Liver | Adult | May 2021 | 3 | 11 | 6 (6) | 6 (6) |
| *C. texanus* | AZ[8,9] | Liver | Adult | May 2021 | 3 | 12 | 6 (6) | 6 (6) |

[1] Colliers Mills Wildlife Management Area, Ocean County, NJ (40°04'N, 74°26′W)
[2] Chiricahua Mountains, Coronado National Forest, Cochise County, AZ (31°54'N, 109°13.1′ W)
[3] Chiricahua Mountains, Coronado National Forest, Cochise County, AZ (31°55.1'N, 109°13.9′ W)
[4] Catalina State Park, Pima County, AZ (32°25.5'N, 110°55′ W)
[5] Rio Vista Natural Resource Park, Pima County, AZ (32°16.8'N, 110°56′W)
[6] Dalquest Desert Research Station, Presidio and Brewster County, TX (29°33.3'N, 103°47.2′W)
[7] Huachuca Mountains, Coronado National Forest, Santa Cruz County, AZ (31°26.5'N, 110°26.4′ W)
[8] Santa Catalina Mountains, Coronado National Forest, Pima County, AZ (32°20.2'N, 110°43.2′W)
[9] Santa Catalina Mountains, Coronado National Forest, Pima County, AZ (32°20.3'N, 110°54.5′W)

**Table S2.** DNA sequencing coverage for individual genes across the 11 chromosomes and unplaced scaffolds in the *Sceloporus undulatus* assembly, reported as average absolute coverage in each sex and as weighted average coverage that corrects per-gene coverage estimates for the mean coverage of each chromosome or scaffold within each individual. The $\log_2$ ratio of the weighted male average (+0.001) to the weighted female average (+0.001) is used to visualize sex differences in DNA coverage and identify potential X-linked and Y-linked regions (separate tabs in this table and colored symbols in Figure S1). Genes with < 10 mean coverage in both sexes are masked from the visualization of these data presented in Figure S1.

[Table S2 is included as separate Excel file].

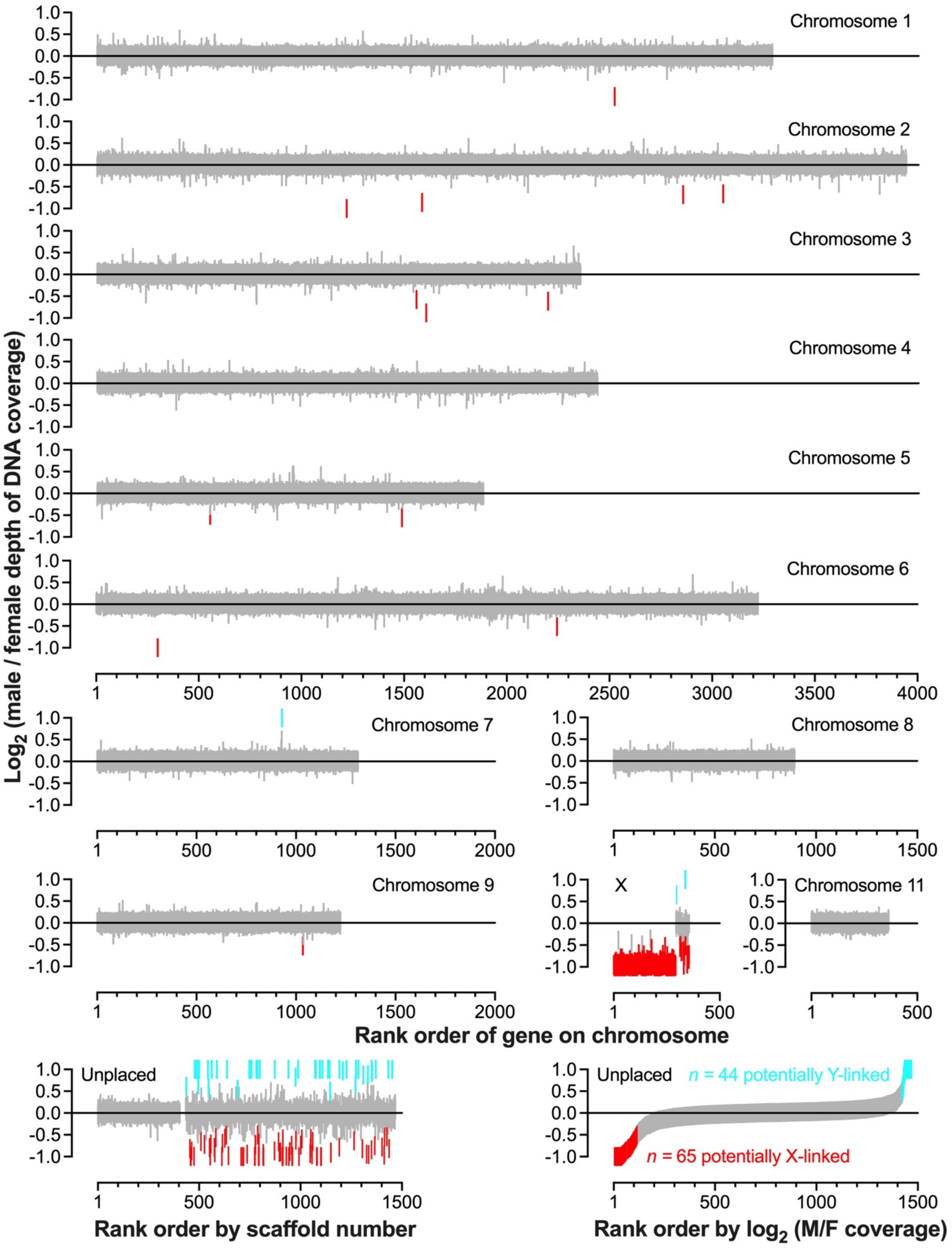
Chromosome 1
Chromosome 2
Chromosome 3
Chromosome 4
Chromosome 5
Chromosome 6
Chromosome 7
Chromosome 8
Chromosome 9
X
Chromosome 11
Unplaced
Unplaced
n = 44 potentially Y-linked
n = 65 potentially X-linked
Log2 (male / female depth of DNA coverage)
Rank order of gene on chromosome
Rank order by scaffold number
Rank order by log2 (M/F coverage)

89

**Figure S1.** Sex differences in DNA sequencing coverage for individual genes across each of the 11 chromosomes (top 11 panels) and aggregated across unplaced scaffolds (lower 2 panels) in the *Sceloporus undulatus* assembly, shown as the $\log_2$ ratio of the weighted male mean ($n$ = 11 individuals) to the weighted female mean ($n$ = 18). Values outside y-axis limits are plotted at -1 or 1 for ease of visualization. Genes with $\log_2$ (male/female) coverage $> 0.5$ are shown in turquoise and those with $\log_2$ (male/female) coverage $< -0.5$ are show in red. Genes with $< 10$ mean absolute depth of coverage in both sexes are masked from this figure (see Table S2). The lower two panels show the same genes on unplaced scaffolds, ranked in two different ways: by scaffold number = scaffold size (left panel), and by $\log_2$ (male/female) coverage (right panel). Nearly all of the $n$ = 109 potentially X- or Y-linked genes highlighted in these two lower panels are located on small scaffolds on which they are the only annotated gene.

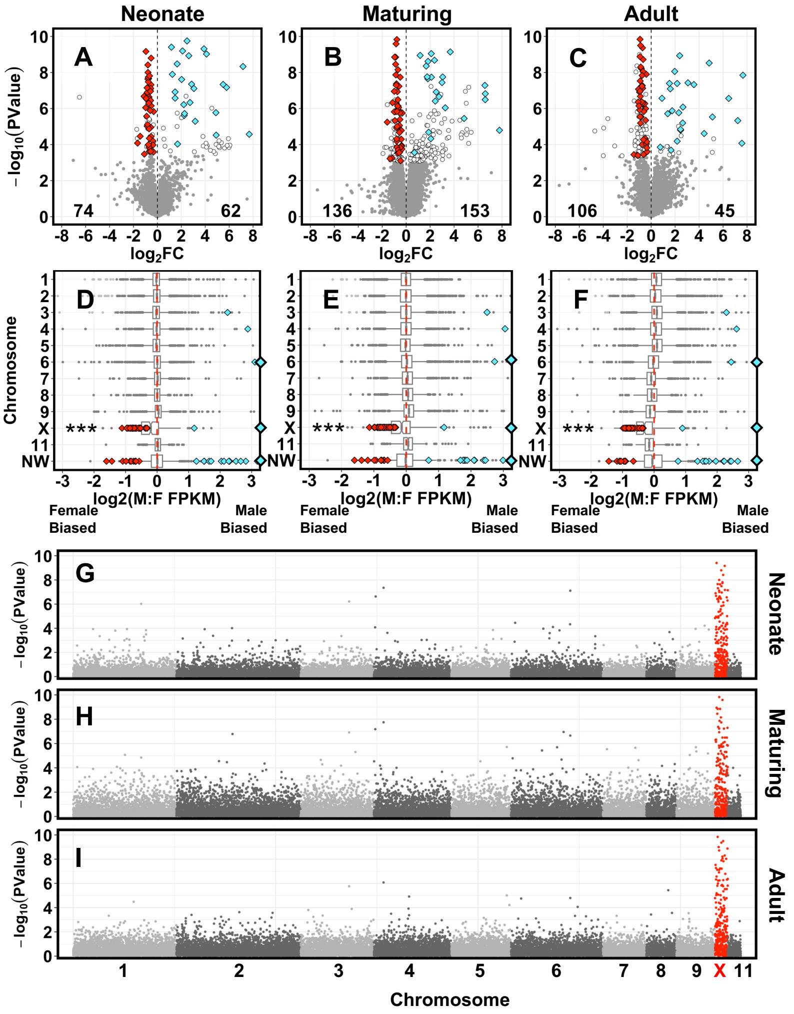


**Figure S2.** Sex-biased expression across 17,423 genes in brain of *Sceloporus undulatus* across three ages, as illustrated with **(A-C)** volcano plots of the $\log_2$ fold change in expression between sexes, **(D-F)** boxplots depicting distributions of sex-biased expression across chromosomes and unplaced scaffolds (NW), and **(G-I)** Manhattan plots of $-\log_{10} P$ values for sex-biased expression after ordering genes by chromosome and start position. Color schemes and statistical tests follow conventions in Figure 2 of the main text, which reports similar patterns in liver.

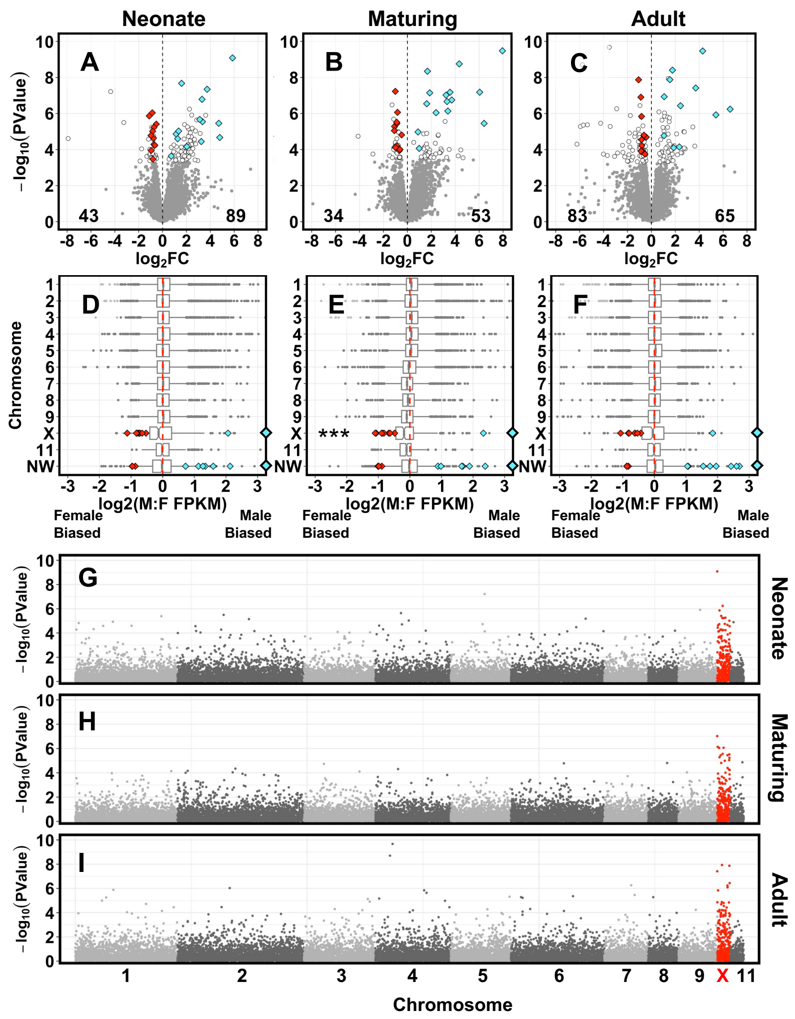


**Figure S3.** Sex-biased expression across 15,135 genes in muscle of *Sceloporus undulatus* across three ages, as illustrated with **(A-C)** volcano plots of the $\log_2$ fold change in expression between sexes, **(D-F)** boxplots depicting distributions of sex-biased expression ($\log_2$(M:F FPKM ratio) across chromosomes and unplaced scaffolds (NW), and **(G-I)** Manhattan plots of $-\log_{10}$ *P* values for sex-biased expression after ordering genes by chromosome and start position. Color schemes and statistical tests follow conventions in Figure 2 of the main text, which reports similar patterns in liver.

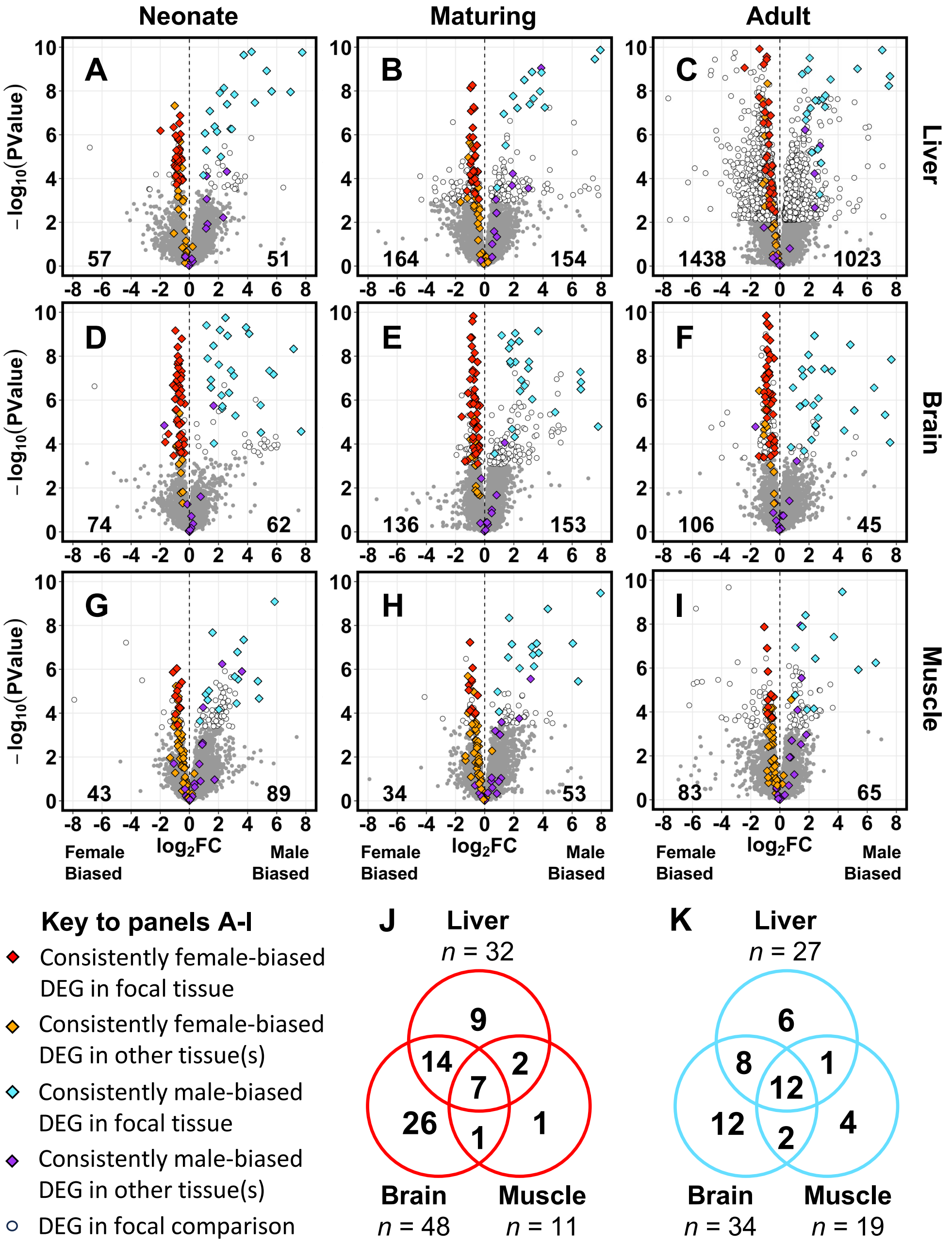


**Figure S4.** Volcano plots depicting differentially expressed genes (DEGs) that are consistently sex-biased across ages in **(A-C)** liver, **(D-F)** brain, and **(G-I)** muscle. Consistently female-biased genes in any given tissue exhibit similar patterns of sex bias across other tissues even if they are not identified as consistently sex-biased DEGs in those tissues. Venn diagrams depict the number of consistently **(J)** female-biased DEGs and **(K)** male-biased DEGs that are shared across tissues.

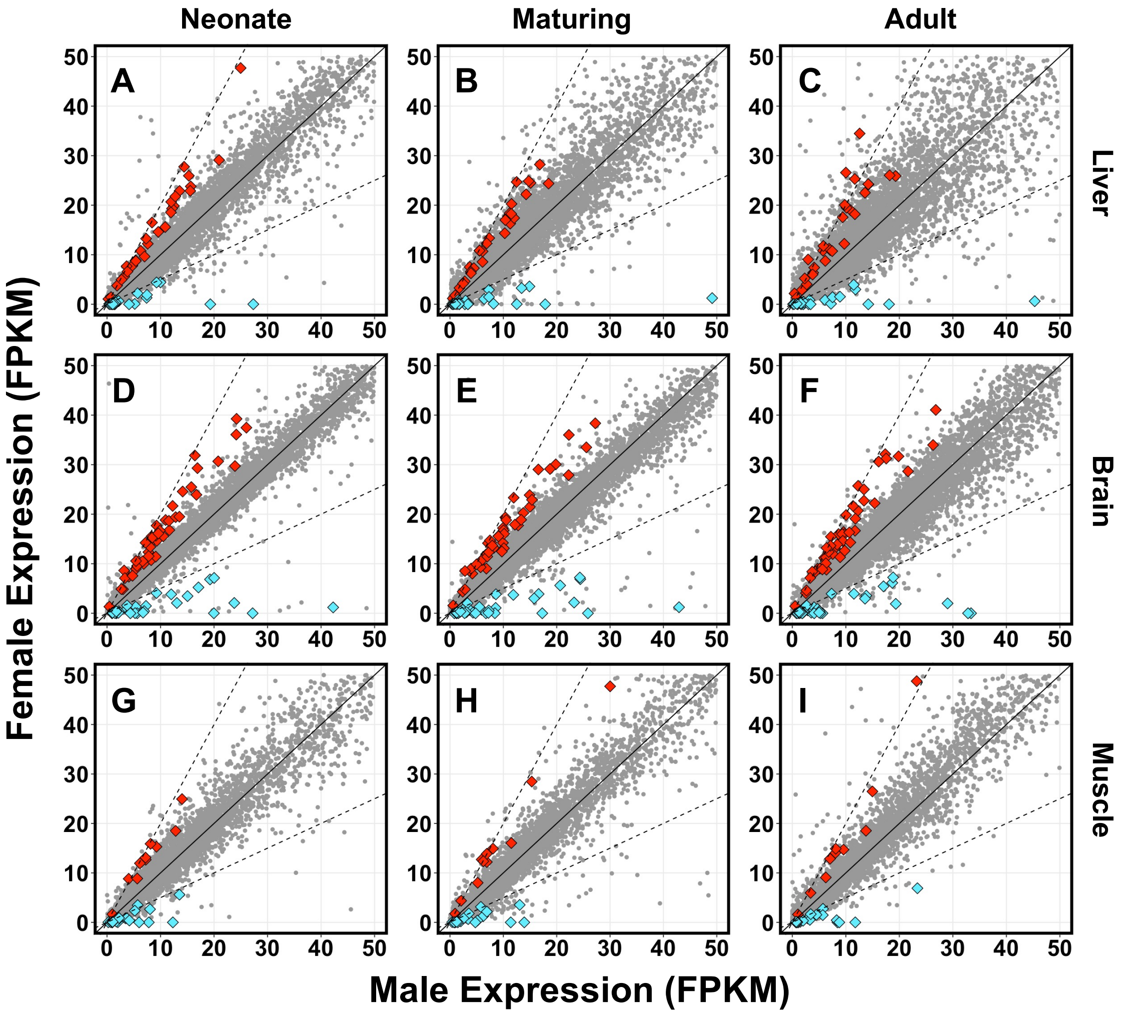


**Figure S5.** Mean expression in females versus mean expression in males (fragments per kilobase per million mapped reads, FPKM) for each gene across three ages in **(A-C)** liver, **(D-F)** brain, and **(G-I)** muscle in *Sceloporus undulatus.* Diamonds indicate consistently female-biased (red) and male-biased (blue) DEGs within each tissue. Gray symbols indicate genes that are not consistently sex-biased within a tissue. Solid diagonal line indicates a 1:1 relationship between male and female expression. Dashed lines indicate 2-fold greater expression in either sex. Axes are truncated at 50 for illustrative purposes.

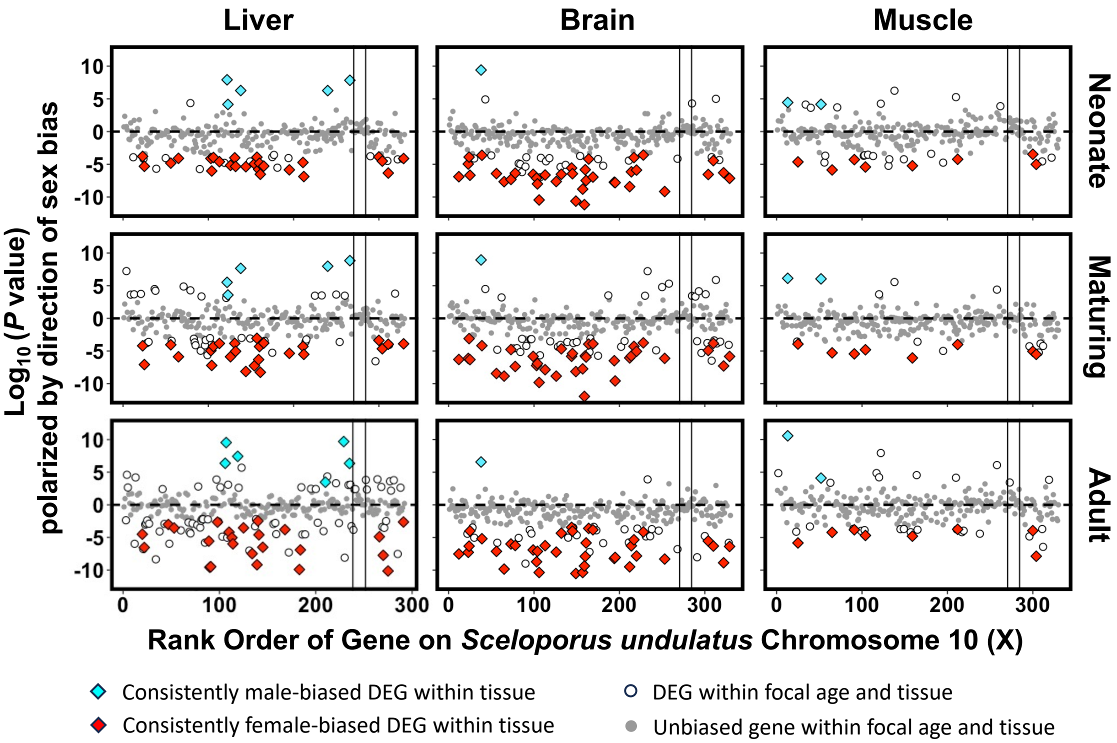


**Figure S6.** Distribution of sex-biased genes across the X chromosome in *Sceloporus undulatus*, as quantified by polarized $-\log_{10}$ (*P* value) from analysis of differential gene expression across ages and tissues. Significantly differentially expressed genes (DEGs) following correction for multiple testing within each combination of age and tissue are indicated with open circles, and genes that are consistently sex-biased within each tissue are indicated with colored diamonds. Unbiased genes are shown in gray. Genes are rank ordered by position along chromosome 10, which is separated by vertical lines into three regions: a proximal 14.85-Mbp region of ancestrally X-linked genes that are hemizygous in males (genes 1-270), an intermediate 0.25-Mbp region of ancestrally X-linked genes with balanced DNA coverage between the sexes that is hypothesized to be a pseudoautosomal region (PAR, genes 271-284), and a distal 1.76-Mbp region that is hypothesized to be more recently X-linked in *S. undulatus* (DR10, genes 285-330). See Figure 1C for corresponding measures of sex differences in DNA coverage for each gene.

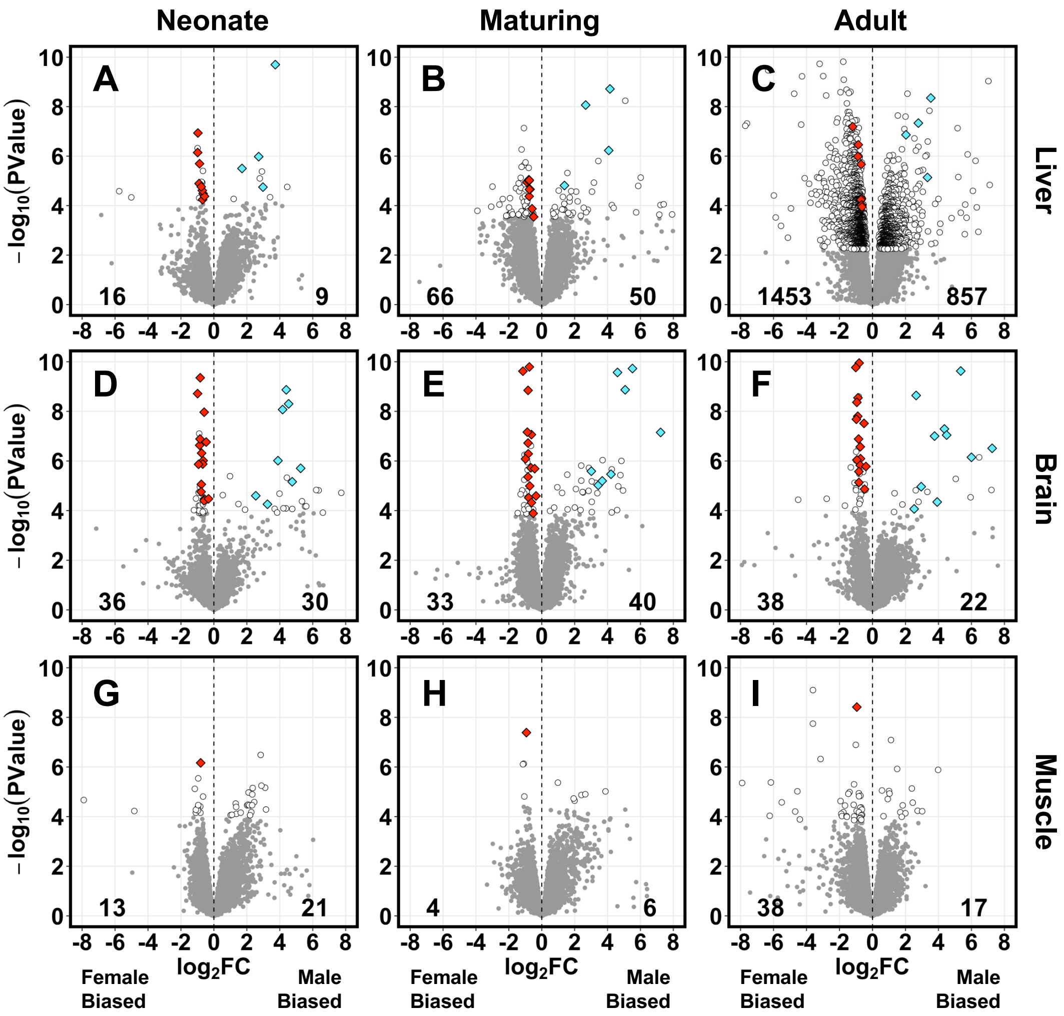


**Figure S7.** Volcano plots depicting the magnitude ($\log_2$ fold change) and significance ($-\log_{10} P$ value) of sex-biased expression in **(A-C)** liver (*n* = 20,288 genes), **(D-F)** brain (*n* = 26,178 genes), and **(G-I)** muscle (*n* = 19,247 genes) of *Sceloporus undulatus* at three ages when mapping reads to the genome assembly of *S. tristichus*, which was sequenced from female tissue and contains no Y chromosome. Diamonds indicate consistently female-biased (red) and male-biased (blue) DEGs within each tissue. White circles indicate DEGs within a given contrast and gray dots indicate unbiased genes. In muscle, sex-biased gene expression was weak and DEGs were identified using an unadjusted $P < 0.01$ threshold; liver and brain DEGs were identified using $P < 0.05$ after adjusting for the false discovery rate.

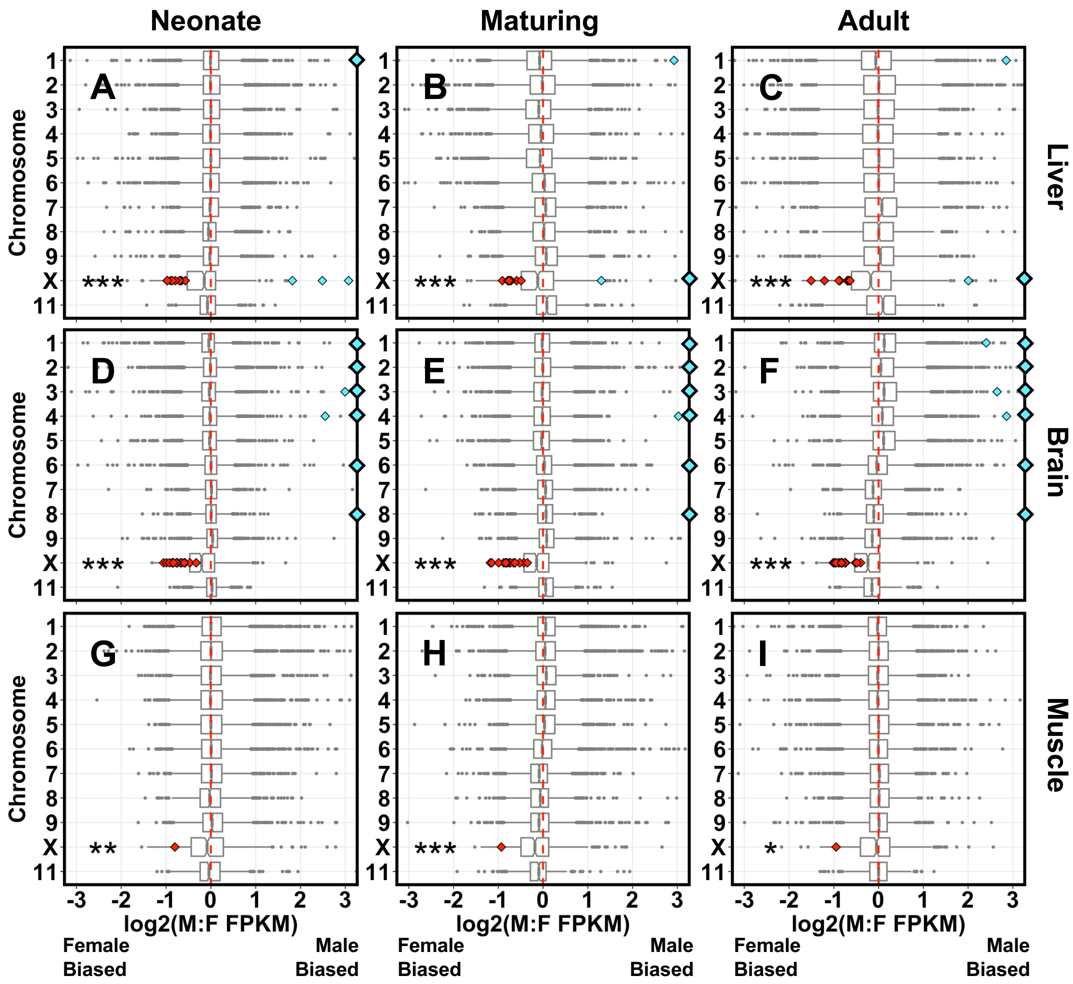


**Figure S8.** Box-and-whisker plots depicting the median (line), interquartile range (box) and the 1.5 x IQR (whiskers) for sex-biased expression ($\log_2$ male:female FPKM) in **(A-C)** liver, **(D-F)** brain, and **(G-I)** muscle of *Sceloporus undulatus* at three ages when mapped to the 11 largest scaffolds in the *S. tristichus* genome. Scaffold 10 is the X chromosome. Notches indicate 95% confidence intervals of boxplot medians. Consistently female-biased DEGs (red diamonds) map to X, while consistently male-biased DEGs (blue diamonds) map to X and autosomes. Diamonds on plot borders indicate genes beyond axis limits. Asterisks indicate significant deviations from balanced male:female expression on X using Wilcoxon signed-rank tests (* $P < 0.05$ without correction; ** $P < 0.017$, the critical alpha after Bonferroni correction for 3 tests; *** $P < 0.001$).

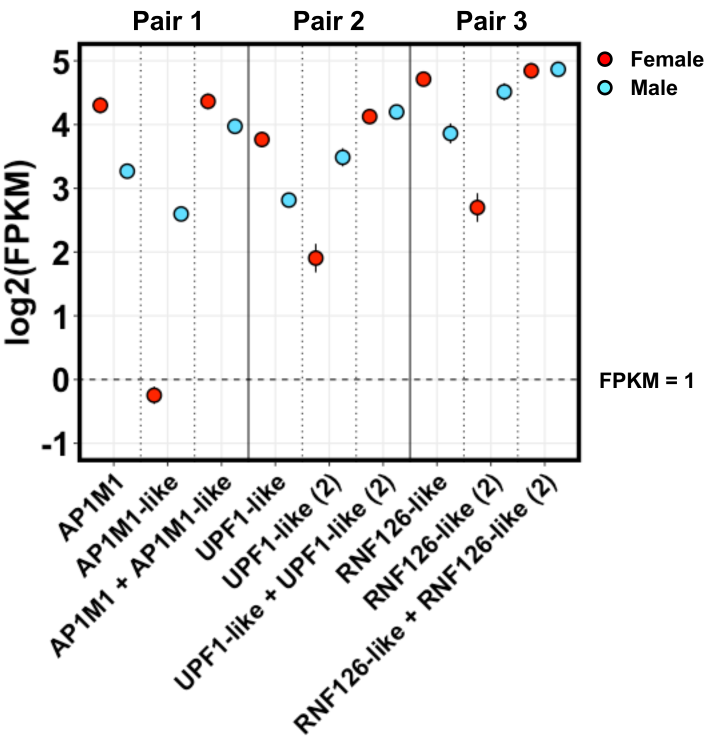


**Figure S9.** Mean (± SEM) expression (log2 FPKM) of three potential gametolog pairs of S. undulatus females (red) and males (blue). Each pair is comprised of one consistently female-biased gene and one consistently male-biased gene, with each mapping to unassembled scaffolds in the S. undulatus genome, and each a reciprocal BLAST hit to the other. The third column for each gene is the summed expression across both genes in the pair (log2(female FPKM + male FPKM). Pair 1, liver gene expression levels: AP1M1 (LOC121917690; consistently female-biased) and AP1M1-like (LOC121917724; consistently male-biased). Pair 2, brain gene expression levels: UPF1-like (LOC121918372; consistently female-biased) and UPF1-like (2) (LOC121918455; consistently male-biased). Pair 3, muscle gene expression levels: RNF126-like (LOC121918298; consistently female-biased) and RNF126-like (2) (LOC121918660; consistently male-biased).

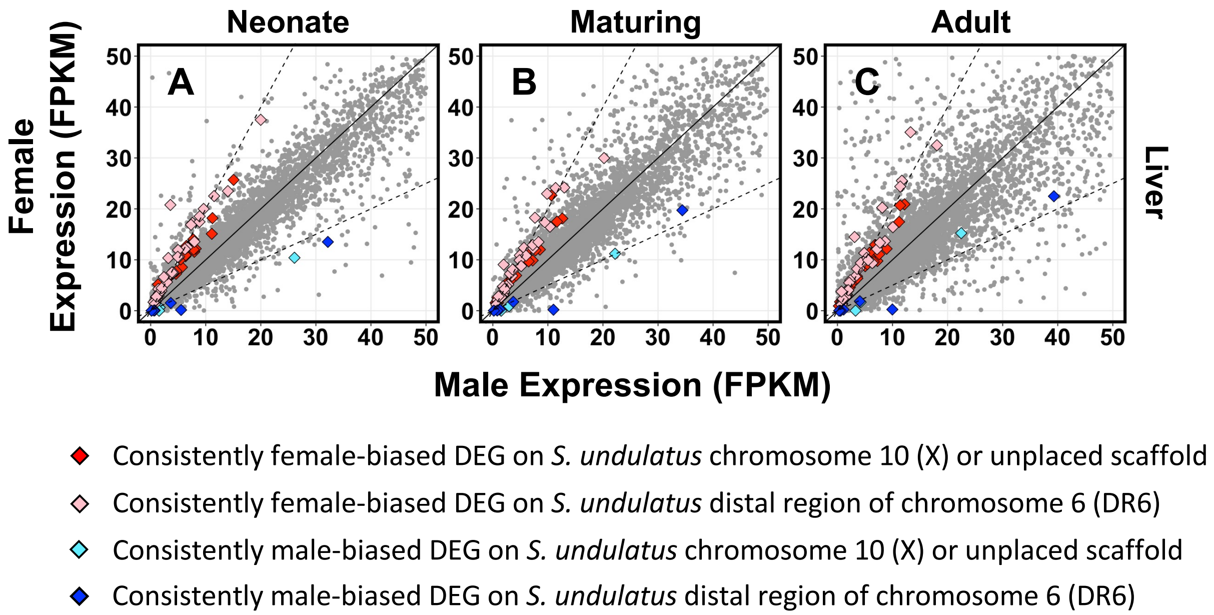


**Figure S10.** Female expression versus male expression (fragments per kilobase per million reads, FPKM) of each gene across three ages **(A-C)** in *Sceloporus jarrovii* liver, based on alignment to the *S. undulatus* assembly. Diamonds indicate genes that are consistently female-biased or male-biased DEGs across all three ages, with the direction of sex bias and chromosomal location in the *S. undulatus* assembly indicated by color in the legend. Genes that are not consistently sex-biased across all three ages are plotted in gray. Solid diagonal line corresponds to balanced 1:1 expression between sexes. Dashed lines correspond to 2-fold greater expression in either sex.

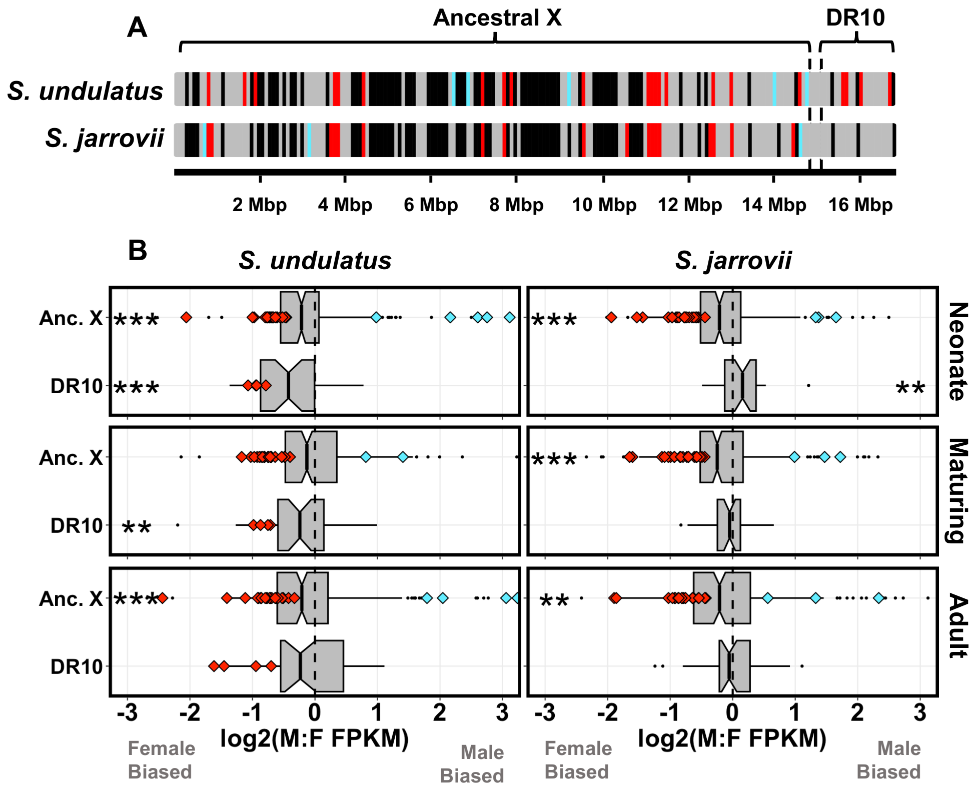


**Figure S11.** (**A**) Location of consistently female-biased (red), male-biased (blue), and unbiased genes (gray) that map to chromosome 10 in *S. undulatus,* plotted separately by expression in *S. undulatus* and *S. jarrovii* liver. Bars are not drawn to scale for gene length. The "Ancestral X" region is hypothesized to be X-linked in both species, while the distal region of chromosome 10 ("DR10") is hypothesized to be X-linked only in *S. undulatus.* (**B**) Boxplots depicting median (line), interquartile range (box), and 1.5 x IQR (whiskers) for sex-biased expression ($\log_2$ male:female ratio of fragments per kilobase per million mapped reads, FPKM) in liver of *S. undulatus* and *S. jarrovii* at three ages. Boxplots are shown separately for the Ancestral X and DR10 regions. Diamonds depict consistently sex-biased DEGs. Asterisks indicate when median expression differs from zero using Wilcoxon signed-rank tests with Bonferroni correction for 2 tests per age (* 0.025 > $P$ > 0.001; ** 0.001 > $P$ > 0.0001; *** $P$ < 0.0001).

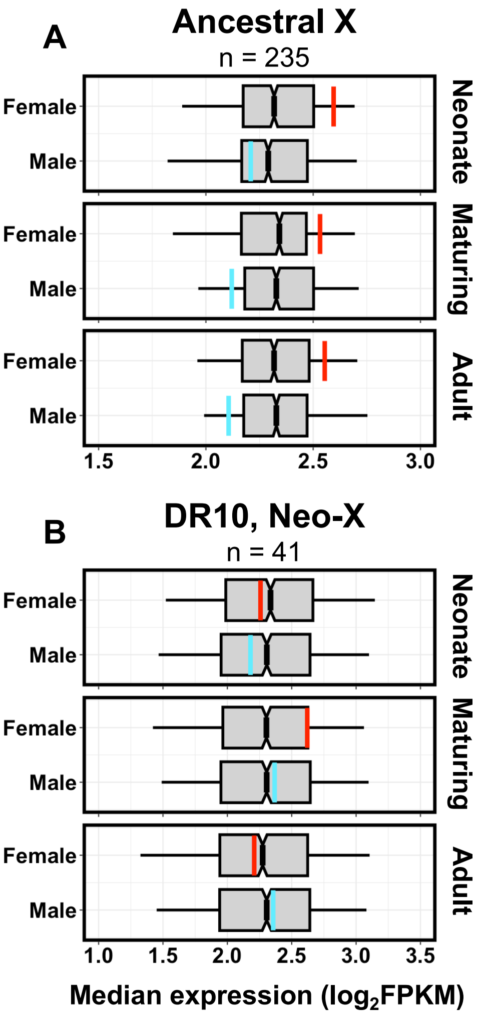


**Figure S12.** Comparisons of median expression across three ages in *S. jarrovii* liver for genes that map to chromosome 10 (X) in the *S. undulatus* assembly. Genes are separated into those that map to **(A)** the proximal 14.85-Mbp region of chromosome 10 ("Ancestral X"), which is hypothesized to be X-linked in *S. jarrovii*, and **(B)** the distal region of chromosome 10 ("DR10"), which is hypothesized to be autosomal in *S. jarrovii.* Boxplots depict the median (line), IQR (box), and 2.5$^{th}$ and 97.5$^{th}$ quantiles (whiskers) for median expression estimates from 1000 randomly sampled blocks of *n* contiguous autosomal genes (*n* = number of expressed genes in the corresponding region of chromosome 10). Colored vertical lines (red = female; blue = male) indicate the median expression of all genes on the corresponding region.
